# LPHN2 inhibits vascular permeability by differential control of endothelial cell adhesion

**DOI:** 10.1101/2020.04.28.065979

**Authors:** Chiara Camillo, Nicola Facchinello, Giulia Villari, Giulia Mana, Noemi Gioelli, Chiara Sandri, Matteo Astone, Dora Tortarolo, Fabiana Clapero, Dafne Gays, Roxana E. Oberkersch, Marco Arese, Luca Tamagnone, Donatella Valdembri, Massimo Santoro, Guido Serini

## Abstract

Dynamic modulation of endothelial cell-to-cell and cell-to-extracellular matrix (ECM) adhesion is essential for blood vessel patterning and functioning. Yet, the molecular mechanisms involved in this process have not been completely deciphered. We identify the adhesion G protein-coupled receptor (ADGR) Latrophilin 2 (LPHN2) as a novel determinant of endothelial cell (EC) adhesion and barrier function. In cultured ECs, endogenous LPHN2 localizes at ECM contacts, signals through cAMP/Rap1, and inhibits focal adhesion (FA) formation and nuclear localization of YAP/TAZ transcriptional regulators, while promoting tight junction (TJ) assembly. ECs also express an endogenous LPHN2 ligand, fibronectin-leucine-rich transmembrane 2 (FLRT2), that prevents ECM-elicited EC behaviors in a LPHN2-dependent manner. Vascular ECs of *lphn2a* knock-out zebrafish embryos become abnormally stretched, display a hyperactive YAP/TAZ pathway, and lack proper intercellular TJs. Consistently, blood vessels are hyperpermeable and intravascularly injected cancer cells extravasate more easily in *lphn2a* null animals. Thus, LPHN2 ligands, such as FLRT2, may be therapeutically exploited to interfere with cancer metastatic dissemination.

**SUMMARY:** Camillo et al. show that the LPHN2 receptor, upon activation by FLRT2 ligand, inhibits focal adhesion formation and promotes tight junction assembly in endothelial cells. Blood vessels of *lphn2a* null animals are hyperpermeable and injected cancer cells extravasate more easily.

## INTRODUCTION

The small GTPase Rap1 drives blood vessel formation and function (Chrzanowska-Wodnicka, 2013) by exerting opposite effects on cell adhesion and motility (Coló et al., 2012; Hong et al., 2007; Lagarrigue et al., 2015, 2016; Lyle et al., 2008), likely due to the involvement of different subcellular pools of Rap1 regulators and effectors (Bos and Pannekoek, 2012). Indeed, the Rap1-GTP-interacting adapter molecule (RIAM) can either promote, through talin, the conformational activation of integrin adhesion receptors at the leading edge of migrating cells (Lagarrigue et al., 2016) or support, *via* the mitogen-activated protein kinase kinase MEK, the disassembly of centrally located integrin-based focal adhesions (FAs) (Coló et al., 2012). In addition, prototypic repulsive guidance cues such as secreted class 3 Semaphorins (SEMA3) control blood vessel patterning (Valdembri et al., 2016; Wälchli et al., 2015) through the cytosolic GTPase activating protein domain of Plexin receptors (Worzfeld et al., 2014) that negatively regulates Rap1 signaling (Gioelli et al., 2018; Wang et al., 2012). On the contrary, G protein-coupled receptors (GPCRs) promote Rap1 GTP loading, *e.g. via* the adenylyl cyclase (AC)/cyclic adenosine monophosphate (cAMP)/exchange protein directly activated by cAMP (EPAC) guanine nucleotide exchange factor cascade (Bos and Pannekoek, 2012; Christensen et al., 2003). Indeed, cAMP/EPAC/Rap1 is required for the formation of extracellular matrix (ECM) mechanosensing FAs that allow ECs to respond to guidance cues (Chrzanowska-Wodnicka, 2017). However, cAMP/EPAC/Rap1 activation by GPCRs can also inhibit EC migration (Avanzato et al., 2016; Hong et al., 2007).

The ability of ECs to modify their behavior in response to ECM patterning and rigidity also relies on the fact that integrin-containing FAs (Karaman and Halder, 2018) signal to suppress the Hippo pathway-dependent inhibitory phosphorylation of the effectors Yes-associated protein (YAP) and WW domain-containing transcription regulator protein 1 (WWTR1) or transcriptional coactivator with PDZ-binding motif (TAZ). In their dephosphorylated active form, YAP and TAZ translocate to the nucleus to control gene transcription (Dupont et al., 2011; Totaro et al., 2018), angiogenic blood vessel formation and function (Choi et al., 2015; Elaimy and Mercurio, 2018; Kim et al., 2017; Nakajima et al., 2017; Neto et al., 2018; Wang et al., 2016, 2017). While tight junctions (TJs) promote Hippo-dependent phosphorylation, inhibition, and cytosolic sequestration of YAP/TAZ, their dephosphorylation, activation, and nuclear translocation, triggered by FAs, involve integrins and their effectors integrin-linked kinase, Src, and focal adhesion kinase (Karaman and Halder, 2018; Moya and Halder, 2019). In addition, myosin-driven contraction of actin stress fibers, which mechanically connect FAs to the nucleus *via* the linker of nucleoskeleton and cytoskeleton (LINC) complex (Kechagia et al., 2019), open nuclear pores to allow YAP/TAZ nuclear translocation (Elosegui-Artola et al., 2017; Kechagia et al., 2019; Totaro et al., 2018).

To identify novel guidance receptors that may be key in the regulation of blood vessel formation and function, we focused on adhesion GPCRs (ADGRs) that interact with several ECM ligands and integrins (Langenhan et al., 2013), whose adhesive functions are regulated by archetypal guidance cues and their receptors, such as SEMA3 and Plexins (Serini et al., 2003; Worzfeld and Offermanns, 2014). In this context, Latrophilin 2 (LPHN2) emerged as an ideal candidate since, similarly to Plexins, it was originally identified as a neuronal receptor (Südhof, 2001) and its mRNA was reported to be highly expressed in *ex vivo* isolated blood vascular, but not lymphatic ECs (Valtcheva et al., 2013). In neurons, the fibronectin leucine-rich transmembrane (FLRT) proteins act as sheddable chemorepulsive ligands (Yamagishi et al., 2011) that bind and signal *via* the LPHN receptors (Seiradake et al., 2016). Here, we unveil how in ECs FLRT2-elicited LPHN2 signals to inhibit ECM adhesion, but to promote TJ assembly, thus limiting YAP/TAZ signaling and vascular permeability.

## RESULTS AND DISCUSSION

To characterize the mechanisms by which LPHN2 receptor controls ECM adhesion, we investigated its subcellular localization and function in cultured human EC. First, we confirmed that endogenous LPHN2 protein is expressed on the EC surface (**Fig. S1A**) and fluorescence confocal microscopy experiments revealed a robust enrichment of endogenous LPHN2 in vinculin containing ECM adhesions of ECs (**Fig. 1A**). Due to autocatalytic processing, ADGRs exist as noncovalently associated heterodimers comprising an extracellular and a transmembrane subunit (Langenhan et al., 2013). To verify the relationships of both LPHN2 subunits with ECM adhesions, we transfected ECs with an N-terminally HA-tagged and C-terminally EGFP-tagged mouse Lphn2 construct (HA-Lphn2-EGFP; **Fig. S1B**). We found that, similarly to endogenous LPHN2, both the HA-tagged extracellular and the GFP-tagged intracellular moieties of HA-Lphn2-EGFP co-localized with vinculin at cell-to-ECM adhesion sites (**Fig. 1B**). Next, we evaluated the outcome of LPHN2 silencing (**Fig. S1C**) on ECM-elicited EC motility (Gioelli et al., 2018). Impedance-based time-lapse migration assays revealed that LPHN2 silenced (siLPHN2) ECs migrate much faster towards type I collagen (Coll I) (**Fig. 1C**) or fibronectin (FN) (**Fig. S1D**) than control silenced (siCTL) ECs. Thus, in cultured ECs, LPHN2 mediates inhibitory signals that are likely initiated by autocrine loops of endogenous LPHN2 ligands.

**Figure 1.**
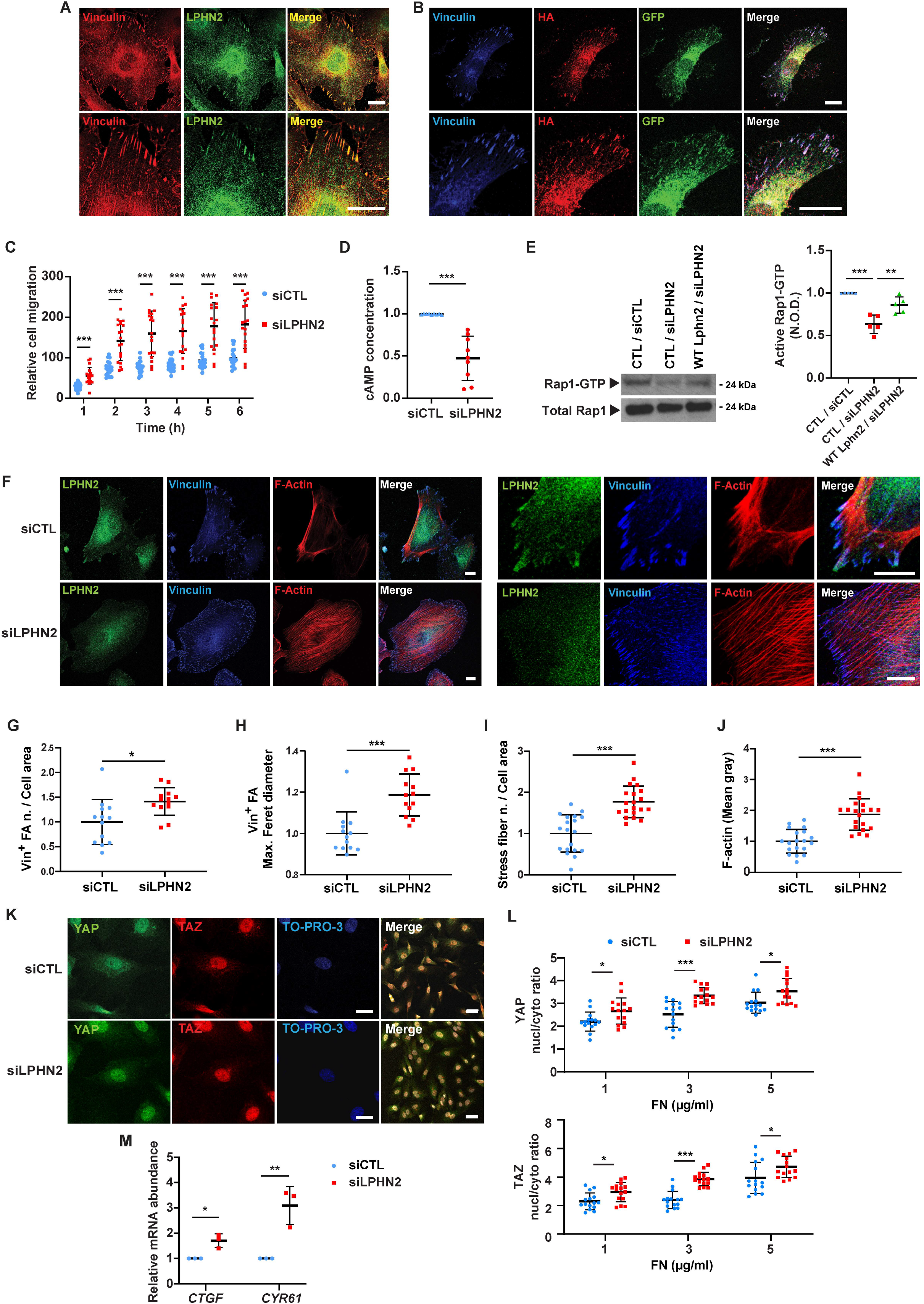
LPHN2 signals *via* cAMP/Rap1 and negatively regulates focal adhesion turnover, stress fiber formation, and ECM-elicited mechanosensing. (**A**) Confocal microscopy analysis of ECs indicates how endogenous LPHN2, as detected by an anti-LPHN2 antibody (*green*), co-localizes with vinculin (*red*) in ECM adhesions of ECs as shown in merge (right panel). Lower row panels are magnifications of the corresponding upper row panels. Scale bars 20 µm. (**B**) Fluorescence confocal microcopy reveals that in ECs transfected with HA-Lphn2-EGFP both the extracellular, as detected by an anti-HA antibody (*red*), and the EGFP-fused intracellular (*green*) moieties co-localize with vinculin (*blue*) at ECM adhesions, as shown in merge (right panel). Lower row panels are magnifications of the corresponding upper row panels. Scale bars 20 µm. (**C**) Real time analysis of cell migration in control (siCTL) or LPHN2 (siLPHN2) silenced ECs toward Coll I, assessed with an xCELLigence RTCA DP system. Results are the average ± SD of five independent assays. Statistical analysis: two-way ANOVA and Bonferroni’s post hoc analysis; p≤ 0,001 ***. (**D**) LPHN2 silencing in ECs decreases basal amount of cAMP. Results are the average ± SD of nine independent assays. Statistical analysis: two-tailed heteroscedastic Student’s t-test; p≤ 0,001 ***. (**E**) Rap1-GTP was pulled down on a glutathione-S-transferase (GST) fusion protein carrying the Rap1-binding domain of human Ral guanine nucleotide dissociation stimulator. LPHN2 silencing in human ECs decreases basal GTP loading of Rap1 small GTPase and pCCL lentivirus-mediated overexpression of silencing-resistant mouse Lphn2 rescues the decrease. Total Rap1 was employed to calculate the normalized optical density (N.O.D.) levels of active Rap1-GTP. Results are the average ± SD of five independent experiments (a representative one is shown). Statistical analysis: one-way ANOVA and Bonferroni’s post hoc analysis; p≤ 0,001 (***) for CTL/siCTL *vs.* CTL/siLPHN2 and p ≤0,01 (**) for CTL/siLPHN2 *vs.* Lphn2/siLPHN2. (**F-J**) Confocal microscopy analysis (**F**) of endogenous LPHN2 (*green*), vinculin (*blue*), and phalloidin-labeled F-actin (*red*; each high magnification images are on the right) reveals how, compared to siCTL ECs, LPHN2 silencing increases the number, normalized on cell area (**G**) and size (expressed by maximum Feret diameter, **H**) of vinculin-containing FAs and the number, normalized on cell area (**I**) and amount (evaluated as mean gray intensity, **J**) of F-actin stress fibers in siLPHN2 ECs. Scale bars 20 µm. Results concerning vinculin-containing FAs and F-actin stress fibers are the average ± SD of two independent experiments for a total of 13 siCTL and 13 siLPHN2 ECs and two independent experiments for a total of 19 siCTL and 20 siLPHN2 ECs, respectively. Statistical analysis: two-tailed heteroscedastic Student’s t-test; p≤ 0,05 *; p ≤0,001 (***). (**K-L**) Confocal microscopy analysis (**K**) of endogenous YAP (*green*) and TAZ (*red*) reveals how, compared to siCTL ECs, LPHN2 silencing increases the nuclear/cytoplasmic ratio of both YAP (**L**, up panel) and TAZ (**L**, bottom panel). Nuclei were stained with TO-PRO3 (*blue*). Scale bar: 20 μm. Representative images of ECs plated on 10kPa stiffness on FN (5 µg/ml) are shown. Results are the average ± SD of 2 independent experiments for a total of 11 ECs for each condition (10kPa and increasing FN 1, 3, and 5 µg/ml concentration). Statistical analysis: two-way ANOVA and Bonferroni’s post hoc analysis; p≤ 0,05 *; p≤ 0,001 ***. (**M**) Real time quantitative PCR analysis of *CTGF* and *CYR61* mRNA in siCTL or siLPHN2 human ECs relative to the house keeping genes, *GAPDH* and *TBP*, and normalized on siCTL levels. Data of one of two independent assays are shown. Results are the average ± SD of three technical replicates. Statistical analysis: two tailed heteroscedastic Student’s t-test; p≤ 0,05 *; p≤ 0,01 **.

Several GPCRs activate the cAMP/Rap1 pathway that regulates cell adhesion (Gloerich and Bos, 2011). Thus, we assessed whether in ECs LPHN2 may impact on basal cAMP production and Rap1 GTP loading by pulling it down with a glutathione-S-transferase (GST) fusion protein carrying the Rap1-binding domain of human Ral guanine nucleotide dissociation stimulator (Franke et al., 1997). Compared to siCTL ECs, siLPHN2 ECs displayed a decrease in the basal amount of both cAMP (**Fig. 1D**) and active Rap1 (**Fig. 1E**) by 52.6 ± 8.7% and 36.35 ± 4.8%, respectively. Moreover, the transduction of a silencing resistant mouse Lphn2 construct (**Fig. S1E**) effectively rescued active Rap1 levels in siLPHN2 ECs (**Fig. 1E**). These findings were consistent with the notion that cAMP/EPAC/Rap1 activation can inhibit migration by modulating cell adhesion dynamics (Lyle et al., 2008). Therefore, we next evaluated the impact of LPHN2 silencing on FAs and the associated F-actin stress fibers. Confocal microscopy analysis of both vinculin (**Fig. 1F**) and paxillin (**Fig. S1F**) containing FAs revealed that in ECs the lack of LPHN2 results in a significant increase of both FA corrected number (**Fig. 1G** and **S1G**) and size (**Fig. 1H** and **S1H**). Moreover, the corrected number (**Fig. 1I**) and mean grey fluorescence (**Fig. 1J**) of F-actin stress fibers were also clearly increased in siLPHN2 compared to siCTL ECs. The small GTPase RhoA controls stress fiber formation and FA turnover (Lawson and Burridge, 2014), and other ADGRs, such as GPR56/ADGRG1, were previously reported to activate RhoA (Paavola and Hall, 2012). Yet, we did not detect any reduction in Rho activation after LPHN2 silencing in ECs (**Fig. S1I**). Hence, LPHN2 activates cAMP/Rap1 signaling and negatively regulates FA and stress fiber formation as well as EC migration towards the ECM in a RhoA-independent manner.

In cultured ECs, LPHN2 silencing results in a significantly increased number of FAs coupled to F-actin filaments (**Fig. 1F, G, I, J** and **Fig. S1F-G**), both structures being part of the mechanically sensitive system that promotes YAP/TAZ translocation into the cell nucleus (Chang et al., 2018; Elosegui-Artola et al., 2017; Karaman and Halder, 2018; Kechagia et al., 2019; Moya and Halder, 2019; Totaro et al., 2018). Of note, endothelial YAP/TAZ signaling controls angiogenesis, vascular barrier maturation, and blood vessel maintenance (Kim et al., 2017; Nakajima et al., 2017; Wang et al., 2017). To investigate whether LPHN2 may impact on YAP/TAZ subcellular localization and signaling, we employed hydrogels with a soft 10 kPa elastic module, previously found to be optimally stiff for cultured EC monolayers (Birukova et al., 2013; Galie et al., 2015; Janmey et al., 2020). In agreement with the notion that ECM coating density is a key determinant of force transmission at integrin-based adhesion sites (Elosegui-Artola et al., 2017; Stanton et al., 2019; Lee et al., 2019), we found that coating 10 kPa hydrogels with increasing amounts of FN resulted in a significantly higher dose-dependent increase of YAP/TAZ nuclear translocation (**Fig. 1K,L**) and transcription of YAP/TAZ target genes *CTGF* and *CYR61* (**Fig. 1M**) in siLPHN2 compared to siCTL ECs. Hence, in ECs, LPHN2 inhibits the formation of FAs and stress fibers along with the ensuing nuclear translocation and transcriptional activity of YAP/TAZ.

Next, we reasoned that the increase of ECM-driven motility (**Fig. 1C** and **Fig. S1D**), FAs (**Fig. 1F-H** and **Fig. S1F-H**), stress fibers (**Fig. 1F,I,J**), and YAP/TAZ nuclear translocation (**Fig. 1K,L**) and signaling (**Fig. 1M**) caused by LPHN2 silencing in ECs may be due to a loss of cell responsivity to endogenously produced LPHN2 ligands, such as FLRT1-3 transmembrane proteins (Seiradake et al., 2016) whose ectodomains can be shed (Jackson et al., 2016) and act *in vivo* as repulsive factors steering both axons (Yamagishi et al., 2011) and blood vessels (Seiradake et al., 2014). In real-time quantitative reverse transcription PCR (qRT-PCR) analyses, we observed that cultured ECs actively transcribe *FLRT2*, but neither *FLRT1* nor *FLRT3* genes (**Fig. S2A**). In addition, we found that the silencing of LPHN2 in ECs causes a two-fold upregulation of *FLRT2* mRNA (**Fig. S2B**), hinting that *FLRT2* and *LPHN2* gene transcription are reciprocally regulated. To explore the potential role of FLRT2 binding in the inhibitory activity of LPHN2 on the formation of FAs and stress fibers, YAP/TAZ accumulation in the nucleus, and the migration rate of ECs, we created a mouse Lphn2 construct devoid of the olfactomedin domain (ΔOLF Lphn2; **Fig. S2C**), which mediates the binding of FLRT ligands to LPHN receptors (Seiradake et al., 2016). Fittingly, ligand-receptor *in situ* binding assay on COS-7 cells demonstrated that FLRT2 interacts with high affinity with wild type (WT), but not ΔOLF Lphn2 (**Fig. S2C**). Next, we compared the abilities of WT and ΔOLF Lphn2 constructs to rescue the aberrant phenotypes of FA corrected number (vinculin^+^ in **Fig. 2A,B** and paxillin^+^ in **Fig. S2 D,E**), size (vinculin^+^ in **Fig. 2A,C** and paxillin^+^ in **Fig. S2D,F**), and stress fibers corrected number (**Fig. 2A,D**), mean grey fluorescence (**Fig. 2A,E**) and mean cross-sectional area (**Fig. 2F**), YAP/TAZ nuclear translocation (**Fig. 2G,H**), and Coll I-elicited migration (**Fig. 2I**) caused by LPHN2 silencing in ECs. Remarkably WT, but not ΔOLF Lphn2 rescued these phenotypic abnormalities in LPHN2 silenced ECs (**Fig. 2** and **Fig. S2D-F**). We concluded that, in ECs, the role played by LPHN2 on the inhibition of FAs, F-actin stress fibers, and ECM-elicited cell motility relies on its OLF domain binding to endogenous ligand(s), such as FLRT2. Consistently, we found that ECM-elicited EC directional migration is respectively increased upon silencing of endogenous FLRT2 (siFLRT2; mRNA in **Fig. S2G** and protein in **Fig. S2H**) (**Fig. 3A**) and decreased by stimulation with exogenous recombinant FLRT2 protein (**Fig. 3B**, left), while no response was seen when LPHN2 was silenced (**Fig. 3B**, right). Similarly to what observed upon LPHN2 silencing (**Fig. 1D,E**), siFLRT2 ECs displayed, compared to siCTL cells, a decrease of the basal amount of both cAMP (**Fig. 3C**) and active Rap1 (**Fig. 3D**) by 12.9 ± 2.9 % and 35.1 ± 6% respectively. In addition, stimulation with exogenous FLRT2 rescued active Rap1 GTP-loading in siFLRT2 ECs (**Fig. 3D**). Furthermore, confocal microscopy revealed that, compared to siCTL, siFLRT2 ECs display a significantly higher corrected number (**Fig. 3E,F**) and size (**Fig. 3E,G**) of vinculin^+^ FAs and F-actin stress fiber corrected number (**Fig. 3E,H**) and mean grey fluorescence (**Fig. 3E,I**). Hence, FLRT2 is one of the endogenous ligands sustaining autocrine/paracrine LPHN2-mediated chemorepulsive signals that negatively regulate the formation of FAs and associated F-actin stress fibers in ECs. Yet, uncleaved FLRT2 may also activate LPHN2 localized outside ECM adhesions, *e.g.* in areas of endothelial cell-to-cell contact.

**Figure 2.**
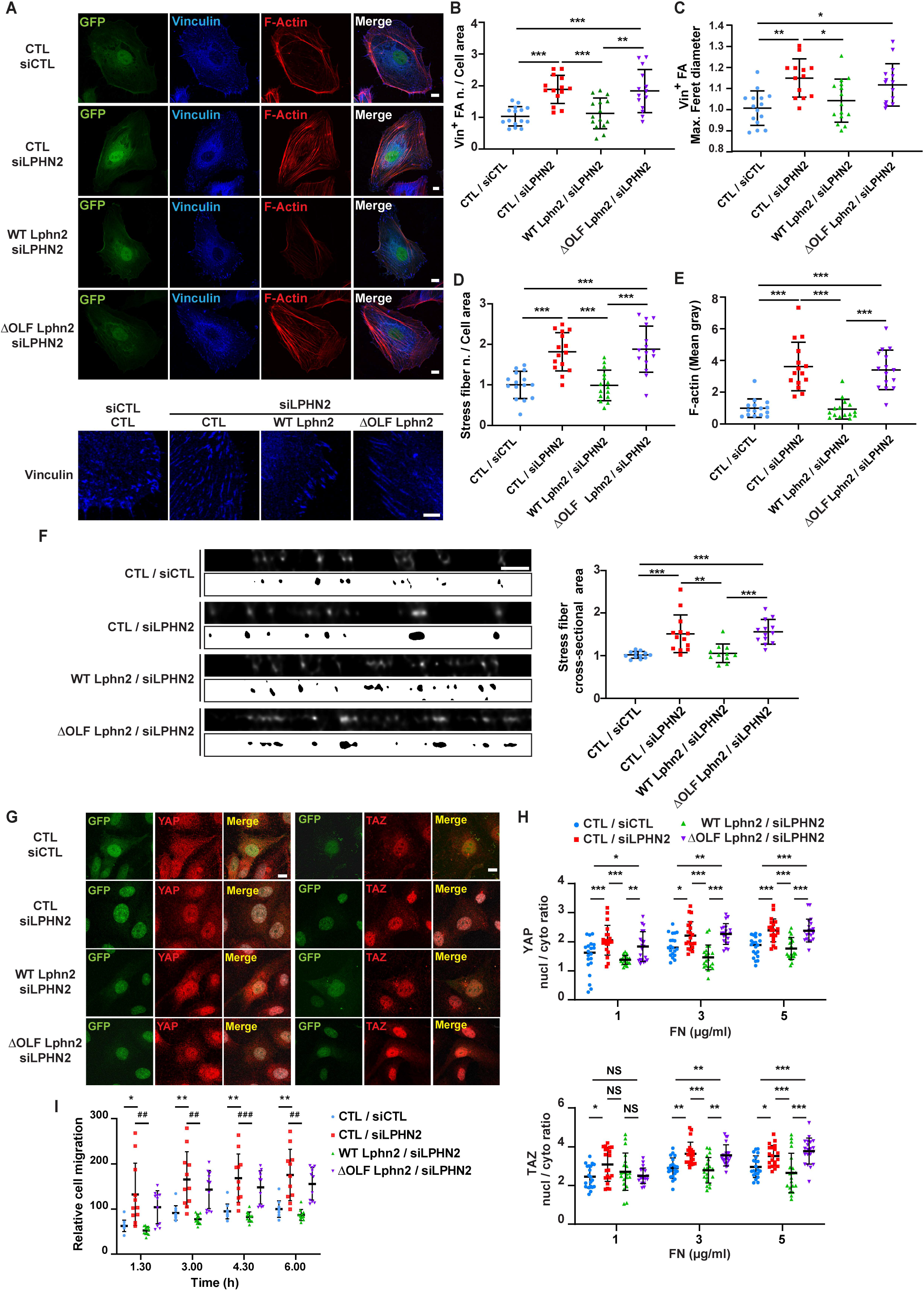
Negative regulation of focal adhesion turnover, stress fiber formation, and ECM-elicited mechanosensing relies on FLRT2-binding OLF domain of LPHN2. (**A-E**) Confocal microscopy analysis (**A**) of vinculin (*blue*), and phalloidin-labeled F-actin (*red*; each vinculin high magnification images are on the bottom). Cells were first transduced with pCCL lentivirus (carrying GFP)-mediated overexpression (*green*) of silencing-resistant mouse WT or ΔOLF Lphn2 and then oligofected with either siCTL or siLPHN2 siRNAs. Scale bars: 10 µm; magnification scale bar: 5 µm. Confocal microscopy analysis reveals how, lentiviral delivery of WT LPHN2, but not ΔOLF LPHN2 mutant, restores the phenotype of vinculin-containing FAs number, normalized on cell area (**B**) and size (expressed by maximum Feret diameter, **C**). The same rescue effect of WT LphnN2, but not ΔOLF Lphn2 mutant occurs on stress fiber number, normalized on cell area (**D**) and amount (evaluated as mean gray intensity, **E**). Results are the average ± SD of 2 independent experiments for a total of 15 ECs for each condition. Statistical analysis: one-way ANOVA and Bonferroni’s post hoc analysis; p≤ 0,05 *; p≤ 0,01 **; p≤ 0,001 ***. (**F**) XZ-STED confocal microscope cross-sectioning of stress fibers reveals how siLPHN2 ECs have thicker stress fibers in comparison to siCTL ECs and the transduction of WT, but not ΔOLF Lphn2, rescues this phenotype. The cross-sectional area of stress fibers was measured with a mask, shown (image with white background) bottom each image, obtained with ImageJ starting from XZ STED confocal images after deconvolution (image with black background). Scale bar: 1 μm. Results are the average ± SD of 2 independent experiments for a total of 11 ECs for each condition. Statistical analysis: one-way ANOVA and Bonferroni’s post hoc analysis; p≤ 0,01 **; p≤ 0,001 ***. (**G-H**) Confocal microscopy analysis (**G**) of endogenous YAP (in *red* on the left) and TAZ (in *red* on the right) reveals how pCCL lentivirus (carrying GFP)-mediated delivery (*green*) of WT Lphn2, but not ΔOLF Lphn2 mutant, restores the nuclear/cytoplasmic ratio of both YAP (**H**, up panel) and TAZ (**H**, bottom panel). Scale bar: 20 μm. Representative images of ECs plated on FN (5 µg/ml)-coated 10 kPa stiff substrate are shown. Results are the average ± SD of 2 independent experiments for a total of 22 ECs for each experimental condition (10kPa and increasing 1, 3, and 5 µg/ml FN concentration). Statistical analysis: two-way ANOVA and Bonferroni’s post hoc analysis; p> 0,05 NS; p≤ 0,05 *; p≤ 0,01 **; p≤ 0,001 ***. (**I**) Real time analysis of EC migration toward Coll I (xCELLigence RTCA DP system) reveals how only transduction with WT (*green*), but not ΔOLF LPHN2 (*purple*) rescues higher migration rate of siLPHN2 ECs. Results are the average ± SD of three independent experiments. Results were analyzed with two-way ANOVA and Bonferroni’s post hoc analysis; p≤ 0,05 *; p≤ 0,01 **.

**Figure 3.**
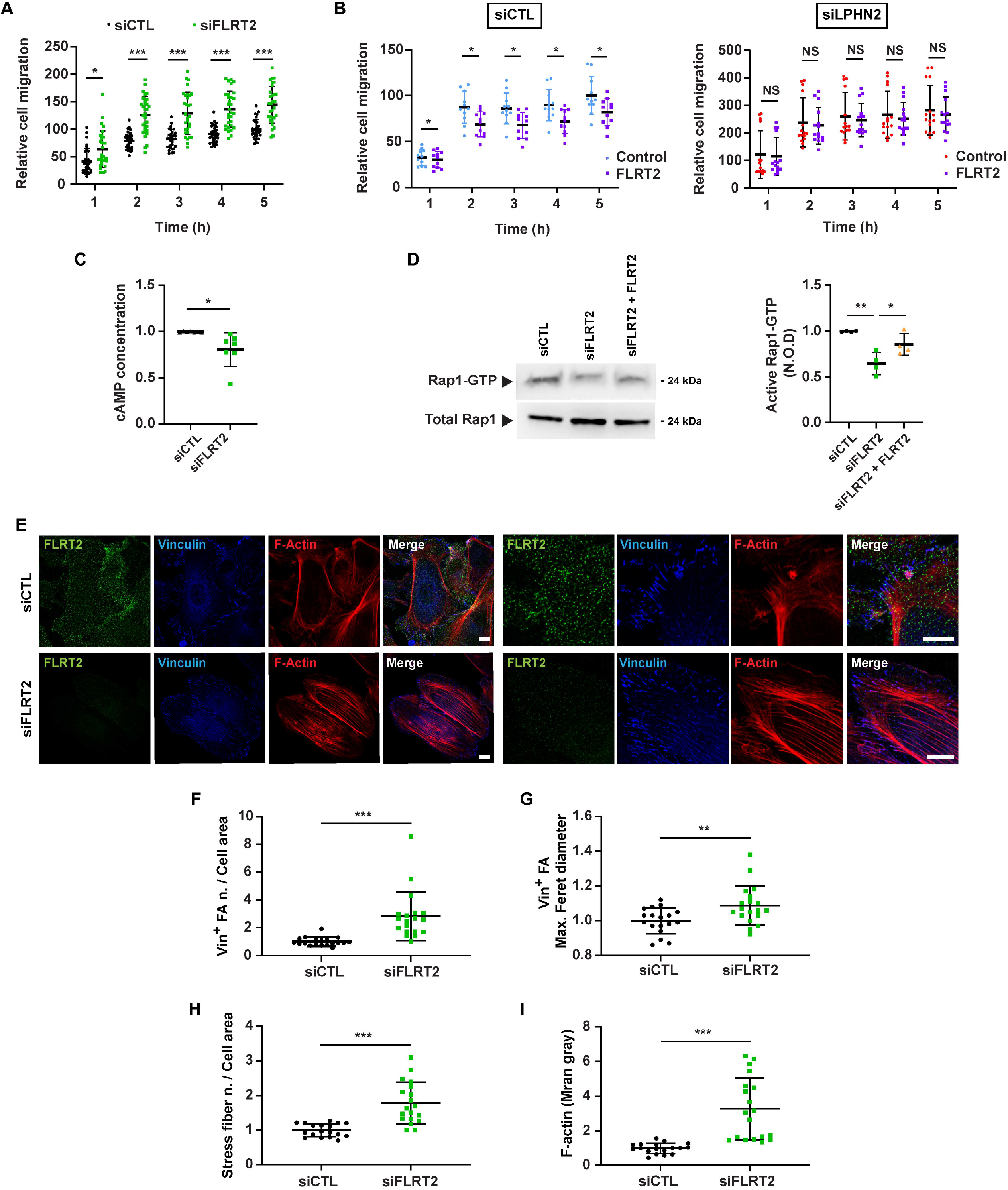
FLRT2 negatively regulates EC response to the ECM *via* LPHN2 and triggers cAMP/Rap1 signaling. **(A, B)** Real time analysis of EC migration toward Coll I, assessed with an xCELLigence RTCA DP system. **(A)** Endogenous FLRT2 silencing (siFLRT2) increases migration compared to siCTL ECs. Results are the average ± SD of seven independent assays. Statistical analysis: two-way ANOVA and Bonferroni’s post hoc analysis; p≤ 0,05 *; p≤ 0,001 ***. **(B)** Exogenous rhFLRT2 [800 ng/ml] inhibits siCTL, but not siLPHN2 EC migration. For sake of simplicity data from the same experiments are plotted in two separated graphs. Results are the average ± SD of three independent assays. Statistical analysis: two-way ANOVA and Bonferroni’s post hoc analysis; p≤ 0,05 * for siCTL ECs (left), whereas all differences in siLPHN2 ECs (right) were not significant, p> 0,05 NS. **(C)** Endogenous FLRT2 silencing in ECs decreases basal cAMP amount. Results are the average ± SD of seven independent assays. Statistical analysis: two-tailed heteroscedastic Student’s t-test; p≤ 0,05 *. **(D)** Endogenous FLRT2 silencing (siFLRT2) in ECs decreases basal Rap1-GTP levels and treatment of siFLRT2 ECs with exogenous rhFLRT2 [800 ng/ml] rescues the decrease in Rap1-GTP levels. Total Rap1 was employed to calculate the normalized optical density (N.O.D.) levels of active Rap1-GTP. Results are the average ± SD of four independent experiments (a representative one is shown). Statistical analysis: one-way ANOVA and Bonferroni’s post hoc analysis; p≤ 0,01 (**) for siCTL *vs.* siFLRT2 and p≤ 0,05 (*) for siFLRT2 *vs*. siFLRT2 + FLRT2. (**E-I**) Confocal microscopy analysis (**E**) of endogenous FLRT2 (*green*), vinculin (*blue*), and phalloidin-labeled F-actin (*red*; each high magnification images are on the right) reveals how, compared to siCTL ECs, FLRT2 silencing increases the number, normalized on cell area (**F**) and size (expressed by maximum Feret diameter, **G**) of vinculin-containing FAs and the number, normalized on cell area (**H**), and amount (evaluated as mean gray intensity, **I**) of F-actin stress fibers in siFLRT2 ECs. Scale bar: 10 µm (siCTL), 15 µm (siFLRT2). Results are the average ± SD of four independent experiments for a total of 18 (siCTL) and 19 (siFLRT2) ECs. Statistical analysis: two-tailed heteroscedastic Student’s t-test; p≤ 0,01 **; p <0,001 ***.

In agreement with what we (**Fig. 1A** and **Fig. S1A**) and others (Valtcheva et al., 2013) observed in cultured ECs, we found that Lphn2a protein is enriched in vascular ECs of dorsal aorta (DA) and posterior cardinal vein (PCV; **Fig. 4A**) of developing transgenic *Tg(kdrl:EGFP)^s843^* zebrafish embryos. Hence, to directly investigate the functional role of LPHN2 in the *in vivo* vasculature, we generated CRISPR/Cas9-mediated *lphn2a* zebrafish knock-out embryos with *Tg(kdrl:EGFP)^s843^* genetic background (**Fig. S3A-D**). The loss of Lphn2a (**Fig. 4A**) did not grossly affect vascular patterning or blood circulation (**Fig. 4B**). In accord with what observed in cultured human ECs (**Fig. S2A-B**), real time qRT-PCR *in vivo* confirmed that *flrt2* gene is actively transcribed in ECs sorted from WT zebrafish embryos and its mRNA levels significantly increase upon *lphn2a* gene knock-down (**Fig. S3E**). *In vitro*, siLPHN2 ECs display a greatly increased number and size of FAs (**Fig. 1F-H; Fig. 2A-C** and **Fig. S1F-H**) coupled to F-actin filaments (**Fig. 1F,I,J** and **2A,D,E**), YAP/TAZ nuclear translocation (**Fig. 1K,L** and **Fig. 2G,H**) and transcription of *CTGF* and *CYR61* target genes (**Fig. 1M**). Therefore, we assessed the impact of *lphn2a* knockdown on the *in vivo* activation of the Hippo pathway in ECs by exploiting the *Tg(Hsa*.*CTGF:nlsmCherry)^ia49^*/ *Tg(kdrl:EGFP)^s843^* double transgenic zebrafish fluorescent reporter line of Yap1/Taz activity (Astone et al., 2018). Of note, *lphn2a* null embryos displayed a significantly increased Yap1/Taz activation in trunk vascular ECs (**Fig. 4C**). Moreover, real-time qRT-PCR revealed a robust upregulation of *ctgfa* and *cyr61* target gene mRNAs in *lphn2a*^-/-^ sorted ECs compared to controls (**Fig. 4D**). Altogether, these *in vivo* findings confirm our *in vitro* model in which FLRT2-activated LPHN2 GPCR inhibits YAP/TAZ signaling.

**Figure 4.**
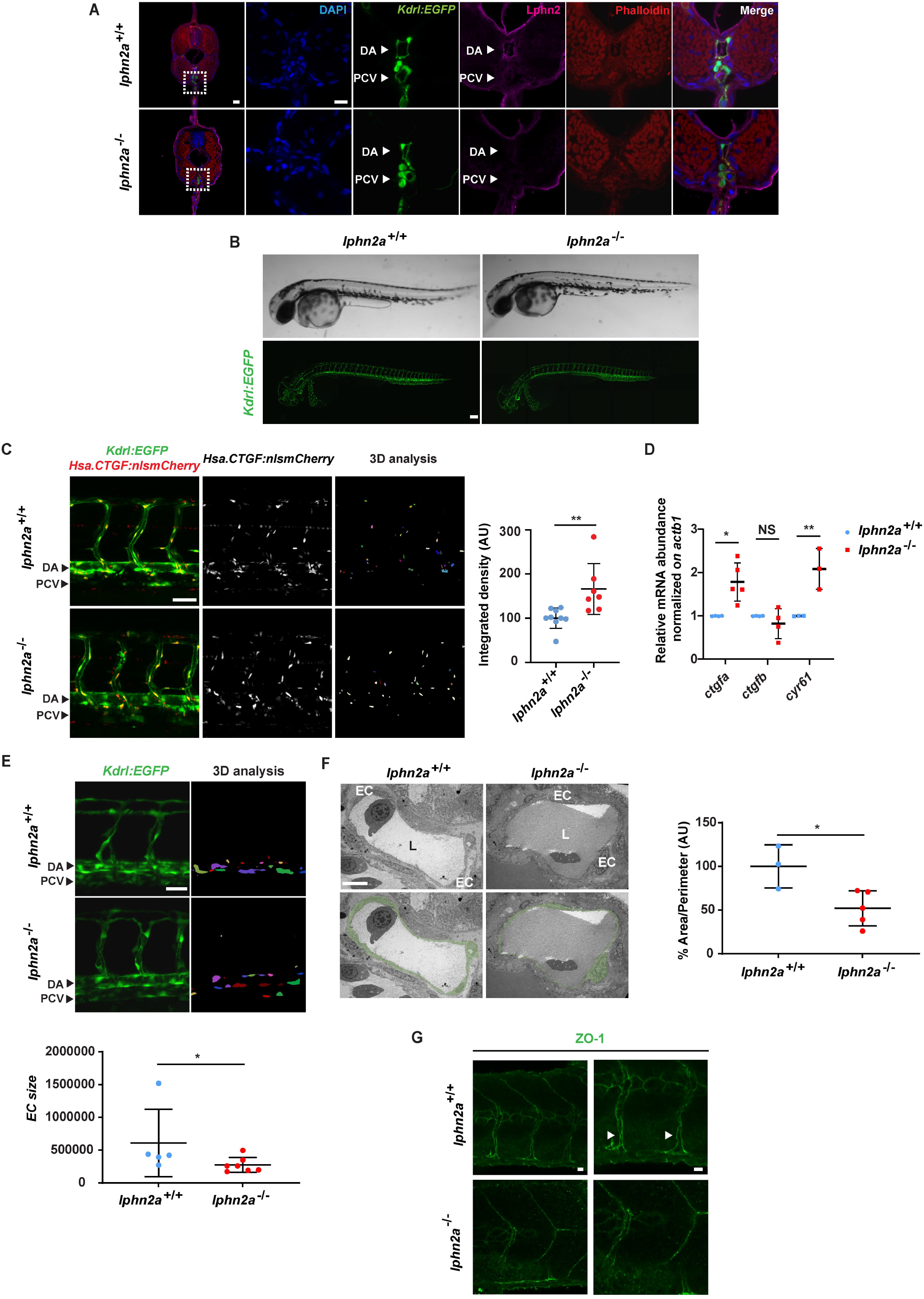
LPHN2 controls EC YAP/TAZ mechanosensing and vascular morphogenesis in zebrafish embryo. (**A**) Confocal fluorescence microscopy analysis of trunk cross-sections at 72 hpf of *Tg*(*Kdrl:EGFP*)*^s843^* WT (*lphn2a*^+/+^) and CRISPR/Cas9-mediated *lphn2a* knock-out (*lphn2a*^-/-^) zebrafish embryos carrying EC-specific EGFP expression (*green*) and stained for Lphn2 (*purple*) and phalloidin (*red*). Nuclei are stained with DAPI (*blue*). Lphn2 is enriched in ECs of dorsal aorta (DA) and posterior cardinal vein (PCV). Whole cross-section in shown in first left panel (scale bar: 30 µm), whereas right one panels display the indicated zoom area (scale bar: 10 µm). (**B**) Lateral view in bright field (up) and fluorescence confocal (bottom) microscopy of WT (*lphn2a*^+/+^) and CRISPR/Cas9-mediated *lphn2a*^-/-^ zebrafish embryos in the *Tg(kdrl:EGFP)^s843^* background. Scale bar: 100 μm. (**C**) Yap1/Taz reporter activity is prominent in the endothelium trunk vasculature of zebrafish embryos. *Left panels*, Representative confocal images of *Tg(Hsa*.*CTGF:nlsmCherry)^ia49^*/*Tg(kdrl:EGFP)^s843^* double transgenic *lphn2a*^+/+^ and *lphn2a*^-/-^ siblings at 60 hpf. Yap1/Taz activation signal was automatically segmented on fluorescent confocal mCherry image Z-stacks, inside a EGFP fluorescent mask identifying endothelial cells. Scale bar: 50 μm. The endothelial cell-restricted Yap1/Taz signal intensity only was represented in 3D analysis and quantified. *Right panel*, Relative quantification of integrated density of Yap1/Taz *Hsa*.*CTGF:nlsmCherry* reporter activity signal co-localized with *kdrl:GFP* in *lphn2a*^+/+^ (n = 9) and *lphn2a*^-/-^ (n = 7) zebrafish embryos (56-72 hpf). Results are the average ± SD. Statistical analysis: Mann-Whitney test; p≤ 0,001 **. (**D**) Real time quantitative PCR analysis of *ctgfa*, *ctgfb*, and *cyr61* mRNAs of FACS-sorted ECs isolated from *lphn2a*^+/+^ or *lphn2a*^-/-^ zebrafish embryos, at 48 hpf, relative to the house keeping gene *actb1* and normalized on the mRNA levels measured in *lphn2a*^+/+^ animals. Results are the average ± SD of ≥3 independent assays (n>80 embryos for condition). Statistical analysis: one-way ANOVA followed by Tukey’s multiple comparison test; p> 0,05 NS; p≤ 0,05 *; p≤ 0,01 **. (**E**) Representative confocal images of *kdrl:GFP* positive cells (left) and represented in the 3D analysis (right) used to quantify EC size (bottom) in *lphn2a^+/+^* (n = 5) and *lphn2a^-/-^* (n = 7). Scale bar: 20 um. Results are the average ± SD. Statistical analysis: Mann-Whitney test; p≤ 0,05 *. (**F**) TEM analysis of endothelium area (left), used to determine the endothelial cell area normalized on the diameter of the vessel (right) in *lphn2a^+/+^* (n = 3) and *lphn2a^-/-^* (n = 5). Color code identifies the EC lumen. Scale bar 10 um. Results are the average ± SD. Statistical analysis: Mann-Whitney test; p≤ 0,05 *. (**G**) Representative confocal images of ZO-1 staining in intersomitic blood vessels (ISVs) of *lphn2a^+/+^* and *lphn2a^-/-^* at 48 hpf. Arrowheads point at continuous ZO-1-stained intercellular contacts between ISV ECs of *lphn2a^+/+^* zebrafish embryos. The ZO-1 intercellular staining between ISV ECs is instead reduced, discontinuous and fragmented in *lphn2a^-/-^* zebrafish embryos. Scale bars: 20 µm (left) and 30 µm (right).

Then, to better define the function of LPHN2 in blood vessels, we thoroughly examined the morphology of vascular ECs of *lphn2a* knock-out *Tg(kdrl:EGFP)^s843^* embryos. Quantitative confocal microscopy showed that ECs forming the trunk vasculature were enlarged, thinner and stretched along the main blood vessel axis in *lphn2a*^-/-^ compared to WT embryos (**Fig. 4E**). Moreover, transmission electron microscopy (TEM) on ultra-thin DA cross-sections of 48 hours post-fertilization (hpf) *lphn2a*^-/-^ zebrafish embryos confirmed that, despite a normal lumen, the lining ECs are extremely thin (**Fig. 4F**). In addition, confocal microscopy revealed a reduced, discontinuous, and fragmented staining of the TJ marker ZO-1 at the intercellular contacts of ECs of intersomitic blood vessels of *lphn2a*^-/-^ embryos compared to WT controls (**Fig. 4G**). Since, in cultured ECs, LPHN2 stimulates the activation of Rap1 (**Fig. 1E**), which promotes TJ formation (Sasaki et al., 2020), LPHN2-elicited Rap1 GTP loading may simultaneously hamper the assembly of FAs and support the developing of TJs. In addition, the recruitment of the mechanosensitive adaptor protein ZO-1 at TJs (Spadaro et al., 2017) was recently described to be under the control of forces regulated by ECM stiffness (Haas et al., 2020). To explore the involvement of LPHN2 in the biochemical and physical crosstalk between FAs and TJs, we plated human ECs on 10 kPa hydrogels coated with increasing FN amounts and assessed, by confocal microscopy, ZO-1 and VE-cadherin (for control purposes) targeting at intercellular contacts. We observed that increasing amounts of FN result in a progressive dose-dependent targeting of ZO-1 at endothelial cell-to-cell contacts in control, but not in LPHN2 silenced ECs (**Fig. 5A**). The VE-cadherin intercellular recruitment was instead not affected by either FN density or LPHN2 silencing (**Fig. 5A** and **Fig. S1L, M**). Furthermore, the lack of FN density-dependent ZO-1 translocation at cell-to-cell contacts in LPHN2 silenced ECs was rescued by the introduction of exogenous WT, but not ΔOLF Lphn2 mutant construct that does not bind FLRT2 (**Fig. 5B**). In sum, LPHN2 controls EC mechanosensing, TJ assembly, and cell shape in living blood vessels. As substantiated in cultured ECs, LPHN2 favors the mechanochemical crosstalk by which FA-sensed ECM stiffness and density promote the formation of TJs.

**Figure 5.**
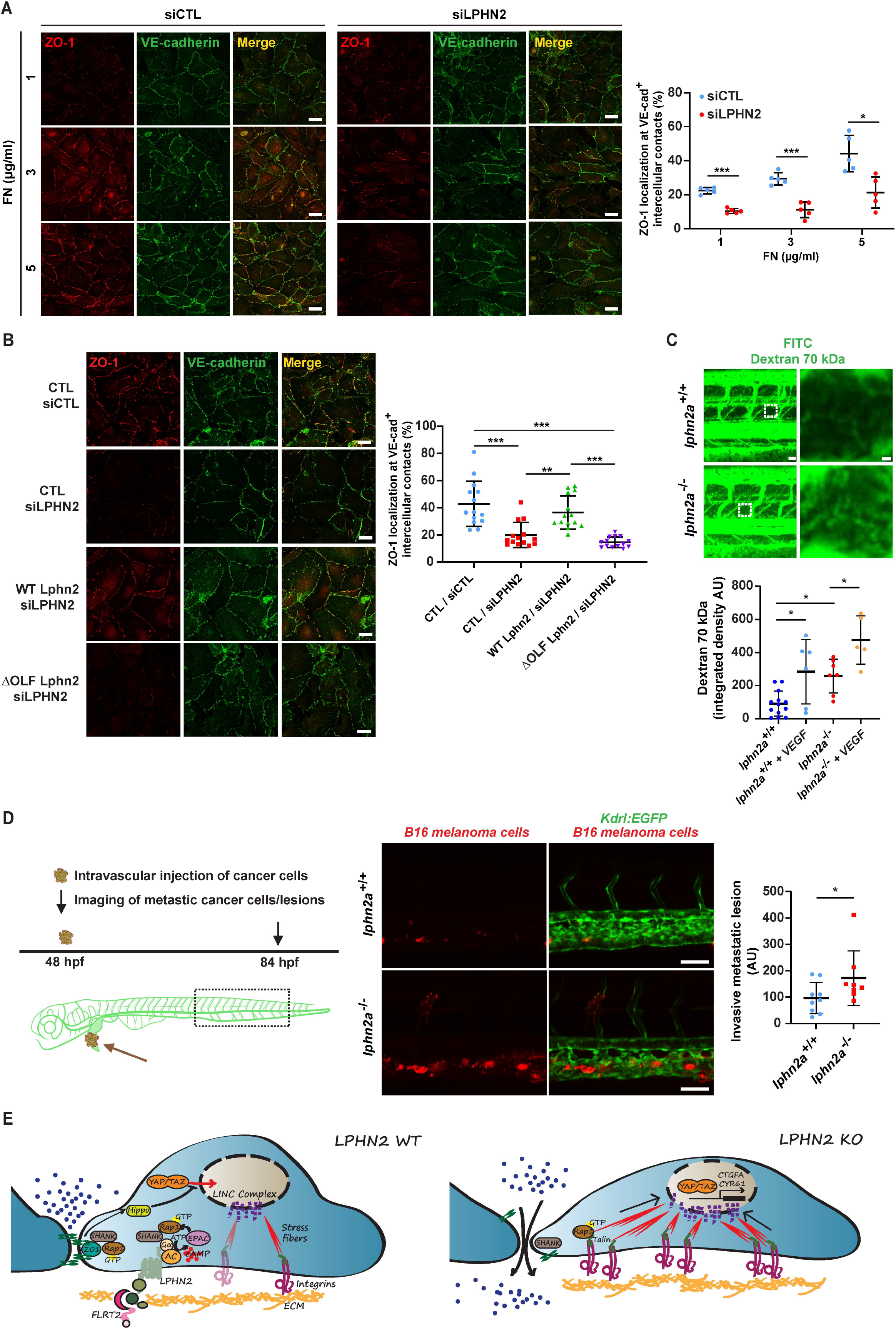
LPHN2 promotes EC tight junction assembly and impairs vascular permeability and cancer cell extravasation. (**A**). Confocal microscopy analysis of TJs stained with ZO-1 (*red*) and VE-cadherin (*green*), reveals how in siCTL ECs seeded on 10 kPa substrates coated with increasing amounts of FN (1, 3, and 5 µg/ml) ZO-1 is progressively accumulating at VE-cadherin^+^ (VE-cad^+^) cell-to-cell junctions. Compared to siCTL ECs, LPHN2 silencing impairs ZO-1, but not VE-cadherin accumulation at intercellular contacts on increasing FN amounts. Scale bar: 25 µm. Results concerning the percentage of VE-cad^+^ intercellular area covered with ZO-1 are the average ± SD of two independent experiments and a total of 5 confocal microscopy images for each condition. Statistical analysis: two-way ANOVA with Bonferroni’s post hoc analysis; p≤ 0,05 *; p≤ 0,001 ***. (**B**). Confocal microscopy analysis reveals how, compared to siCTL ECs, LPHN2 silencing impairs ZO-1 accumulation to VE-cad^+^ intercellular contacts of ECs plated on FN (5 µg/ml)-coated coverslips. Lentiviral delivery of WT Lphn2 restores ZO1 localization to TJs, while the ΔOLF Lphn2 mutant does not rescue the phenotype. Scale bar: 25 µm. Results concerning the percentage of VE-cad^+^ intercellular area covered with ZO-1 are the average ± SD of three independent experiments for a total of 14 confocal microscopy images for each condition. Statistical analysis: one-way ANOVA and Bonferroni’s post hoc analysis; p≤ 0,05 **; p≤ 0,001 ***. (**C**) Up panels, representative images of vascular permeability in *lphn2a*^+/+^ *vs lphn2a*^-/-^ zebrafish embryos. 70 kDa FITC-dextran was injected with or without 1 ng of VEGF-A (Sigma #V7259). 70 kDa Dextran is in *green*. Scale bars: 30 μm (left) and 3 μm (right). Bottom panel, quantification of relative extravascular fluorescence. For each embryo, the fluorescence intensity of the dextran was measured in two intervascular areas between the intersegmental vessels (dashed box and shown in the zoom images on the right). *lphn2a^+/+^* (n = 13), *lphn2a^+/+^* with VEGF (n = 6), *lphn2a^-/-^* (n = 7), *lphn2a^-/-^* with VEGF (n=5) embryos from 2 independent experiments. Results are the average ± SD. Statistical analysis: one-way ANOVA followed by Tukey’s multiple comparison test; p≤ 0,05 *. (**D**) MemBright-560-labeled mouse B16F10 melanoma cells were microinjected into the duct of Cuvier of 48 hpf *lphn2a*^+/+^ or *lphn2a*^-/-^ *Tg*(*Kdrl:EGFP*)*^s843^* zebrafish embryos. After 36 hours post-injection, extravasated metastatic melanoma cells were imaged by confocal analysis of the caudal plexus. Mouse B16F10 melanoma cell extravasation is enhanced in *lphn2a*^-/-^ compared to *lphn2a*^+/+^ zebrafish embryos. Results are the average ± SD of 2 independent assays, in which 17 animals were analyzed. Scale bars: 50 μm. Statistical analysis: Mann-Whitney test; p≤ 0,05 *. (**E**) LPHN2 signaling controls endothelial FAs, TJs, and vascular permeability. Vascular ECs synthesize the FLRT2 ligand of LPHN2 that localizes at integrin-based ECM adhesion sites. FLRT2-activated LPHN2 triggers a canonical heterotrimeric G-protein α subunit (Gα)/adenylate cyclase (AC)/ cAMP pathway that in turn, likely *via* the guanine nucleotide exchange factor EPAC, activates the small GTPase Rap1, which is a well-known regulator of cell-to-ECM adhesions (Coló et al., 2012; Lagarrigue et al., 2016). Furthermore, Rap1 promotes the formation of TJs (Sasaki et al., 2020), which in ECs are crucial for the control of vascular permeability. Hence, LPHN2 activation of Rap1 may act both to inhibit the formation of FAs and to promote the assembly of TJs, which increase EC barrier function. LPHN2 also binds the central PDZ domain of SHANK adaptor protein that in turn through its N-terminal SPN domain binds Rap1-GTP, suppressing talin-mediated integrin activation and FA development (Lilja et al., 2017) and promoting the assembly of TJs (Sasaki et al., 2020). Therefore, LPHN2 may favor the turnover of FAs and the formation of TJs by funneling Rap1-GTP towards SHANK. In addition, while TJs inhibit the nuclear translocation of YAP and TAZ through their Hippo pathway-dependent phosphorylation, FAs and the associated F-actin stress fibers exert exactly the opposite effect promoting YAP/TAZ nuclear localization and transcriptional function (Karaman and Halder, 2018; Moya and Halder, 2019). Thus, the nuclear translocation and functional activation of YAP/TAZ caused by LPHN2 silencing or knock down likely lie downstream of both the disassembly of TJs and the increased formation of FAs and stress fibers. In addition, the myosin II-mediated contraction of FA-linked stress fibers releases YAP/TAZ from their binding to the inhibitory SWI/SNF complex (not depicted) and transmits force from the ECM to the nucleus, changing nuclear pore conformation, finally promoting the translocation of YAP/TAZ into the nucleus and the transcription of target genes, such as *CTGFA* and *CYR61*. The lack of LPHN2 also results in an abnormal ECM-driven intercellular targeting of ZO-1 and assembly of TJs, which increases vascular permeability and favors cancer cell extravasation.

The entry and exit of solutes, leukocytes, and cancer cells from the bloodstream is regulated by the endothelial barrier that relies on the controlled remodeling of TJs (Wettschureck et al., 2019; Zihni et al., 2016). Since we found that the lack of LPHN2 results in the abnormal organization of TJs between ECs *in vivo* (**Fig. 4G,H**) and *in vitro* (**Fig. 5A,B**), we sought to investigate the impact of LPHN2 on both basal and VEGF-A-elicited blood vessel permeability. First, we injected or not *lphn2a*^+/+^ or *lphn2a*^-/-^ zebrafish embryos with VEGF-A. Next, we measured vascular leakage by intravascularly inoculating and measuring the amount of extravasated FITC-Dextran 70 kDa. Both in basal conditions and upon VEGF-A stimulation, FITC-Dextran 70 kDa extravasated significantly more in *lphn2a*^-/-^ than in *lphn2a*^+/+^ zebrafish embryos (**Fig. 5C**). Then, we measured the extravasation of cancer cells that can be effectively tracked and quantified by using the zebrafish embryo (Follain et al., 2018; Osmani and Goetz, 2019). To this aim, fluorescently-labeled mouse (B16F10) or human (SK-MEL-28) melanoma cells were injected in the duct of Cuvier of *Tg(kdrl:EGFP)^s843^* WT or *lphn2a*^-/-^ zebrafish embryos and their metastatic extravasation potential was quantified by measuring the number of tumor cells found outside the vessels and in the caudal plexus where circulating tumor cells (CTCs) preferentially arrest and extravasate (Follain et al., 2018; Hyenne et al., 2019). We discovered that, likely due to their abnormal inter-EC TJs, *lphn2a* null zebrafish embryos are more permissive to extravasation of both mouse B16F10 (**Fig. 5D**) and human SK-MEL-28 (**Fig. S3F**) melanoma cells compared to WT embryos.

To conclude, we identified the endothelial ADGR LPHN2 as a novel repulsive guidance receptor that controls *in vivo* blood vessel structure and function. Our data suggest that, upon activation by ligands, such as FLRT2, LPHN2 elicits the synthesis of cAMP and Rap1-GTP pools that, in turn, promote the disassembly of integrin-based FAs and the assembly ZO-1 containing TJs (**Fig. 5E**). All LPHNs contain a C-terminal motif that specifically binds the PSD-95/discs large/ZO-1 (PDZ) domain of SH3 and multiple ankyrin repeat domains (SHANK) scaffold proteins (Kreienkamp et al., 2000), which were found to compete with talin for binding to Rap1-GTP and impair Rap1-GTP/talin-driven integrin activation (Lilja et al., 2017). Of note, we observed that LPHN2 silencing significantly reduces the physical interaction between Rap1 small GTPase and the scaffold protein SHANK2 in cultured ECs (**Fig. S2I**). Hence, LPHN2 may promote FA turnover by locally funneling Rap1-GTP towards SHANK rather than talin. Interestingly, Rap1-GTP-bound SHANK2 was recently reported to support TJ formation in epithelial cells (Sasaki et al., 2020). Although further work is required, it is tempting to speculate that in ECs LPHN2 may couple FA disruption and TJ formation by promoting Rap1 GTP loading and its binding to SHANK proteins rather than with talin.

We also uncovered that LPHN2 signals by inhibiting the Hippo effectors YAP and TAZ in vascular ECs, both *in vitro* (**Fig. 1K, L**) and *in vivo* (**Fig. 4C, D**). Although other GPCRs were described to modulate YAP/TAZ activation *via* Rho GTPase-regulated F-actin dynamics (Totaro et al., 2018; Yu et al., 2015), we did not detect any effect of LPHN2 silencing on Rho activation (**Fig. S1I**). Hence, the inhibition of the formation of FAs and F-actin stress fibers along with the stimulation of TJ assembly that we report here appear as the most conceivable mechanisms by which LPHN2 negatively regulates YAP/TAZ signaling in ECs (**Fig. 5E**). Indeed, while TJs promote their Hippo-dependent inhibition and cytosolic sequestration, FAs stimulates the activation and nuclear translocation of YAP and TAZ (Karaman and Halder, 2018; Moya and Halder, 2019). Finally, our data show that, likely due to the loosening of endothelial TJs, the lack of Lphn2a favors vascular permeability and the extravasation of tumor melanoma cells (**Fig. 5E**). Such findings indicate that LPHN2 ligands, *e.g.* FLRT2, might be therapeutically exploited to strengthen the vascular barrier and to counteract cancer cell metastatic dissemination.

## MATERIALS AND METHODS

### DNA constructs

Mouse hemagglutinin (HA) tagged Lphn2 construct was synthesized by GeneArt Gene Synthesis (Thermo Fisher Scientific) and cloned into the Gateway system pENTR221 vector (Thermo Fisher Scientific). HA-Lphn2 cDNA was then subcloned into the pEGFP-N1 [Enhanced Green Fluorescent Protein (EGFP)] mammalian expression vector by standard PCR protocols. HA-Lphn2 ΔOLF pEGFP-N1 mutant construct lacking the Olfactomedin domain (amino acids 142-398) was obtained by PCR, according to the Taq polymerase manufacturer’s instructions, using Phusion Site-Directed Mutagenesis Kit (Thermo Fisher Scientific). WT and ΔOLF HA-Lphn2 were also subcloned into the pCCL.sin.cPPT.polyA.CTE.eGFP.minhCMV.hPGK.Wpre (pCCL) lentiviral vector. Lentiviral particles were produced as previously described (Gagliardi et al., 2012). Briefly, transduction of cells was performed with a multiplicity of infection equal to 3 in the presence of 8 µg/ml Polybrene (H-9268; Sigma-Aldrich). Cells were then selected with 2.5 µg/ml puromycin for 2 days, and the surviving cell population was used for the experiments.

### Cell culture

Primary human ECs were isolated from the umbilical cords as previously described (Jaffe et al., 1973). Briefly, umbilical vein was cannulated with a blunt 17-gauge needle that was secured by clamping. The umbilical vein was then perfused with 50 ml of phosphate-buffered saline (PBS) to wash out the blood. Next, 10 ml of 0.2% collagenase A (Cat. # 11088793001, Roche Diagnostics) diluted in cell culture medium were infused into the umbilical vein and incubated 30 min at room temperature. The collagenase solution containing the ECs was flushed from the cord by perfusion with 40 ml of PBS, collected in a sterile 50 ml centrifuge tube, and centrifuged 5 min at 800 *g*. Cells were first resuspended in M199 medium completed with cow brain extract, heparin sodium salt from porcine intestinal mucosa [0,025 mg/500 ml], penicillin/streptomycin solution, 20% fetal bovine serum (FBS) (Sigma Aldrich), and subsequently plated in cell culture dishes that had been previously adsorbed with 1% gelatin from porcine skin (G9136, Sigma Aldrich). Cells were tested for mycoplasma contamination by means of Venor GeM Mycoplasma Detection Kit (MP0025-1KT, Sigma Aldrich) and grown in M199 complete medium. The isolation of primary venous ECs from human umbilical cords was approved by the Office of the General Director and Ethics Committee of the Azienda Sanitaria Ospedaliera Ordine Mauriziano di Torino hospital (protocol approval no. 586, Oct 22 2012 and no. 26884, Aug 28 2014) and informed consent was obtained from each patient. COS-7, HEK 293T, and B16F10 cells (ATCC) were grown in DMEM medium completed with glutamine, penicillin/streptomycin solution, and 10% FBS (Sigma Aldrich). SK-MEL-28 (ATCC) were instead grown in EMEM medium completed with glutamine, penicillin/streptomycin solution, and 10% FBS (Sigma Aldrich). Both COS-7 and HEK 293T were transfected by means of Lipofectamine and PLUS reagent (Thermo Fisher Scientific). The calcium phosphate transfection method was instead used to transfect WT or ΔOLF Lphn2-HA-pCCL in HEK 293T cells and to produce the corresponding viral particles subsequently employed to transduce ECs. To measure cell signaling response to matrix stiffness, ECs were plated on fibronectin-coated PAA gels [6% (∼10 kPa) of gel diluted from 30% Protogel in PBS, 37.5:1 fixed ratio of acrylamide:bis-acrylamide; EC-890, National Diagnostics], previously prepared in glass slide with removable silicon chamber (IBIDI) according to published methods, with modifications (Wang and Pelham, 1998; Zhang et al., 2013).

### Antibodies and reagents

2Y4824 rabbit polyclonal antibody (pAb) anti-LPHN2 was produced by Eurogentec by immunizing animals with peptide GGKTDIDLAVDENGL (amino acids 259-274) (Herberth et al., 2005). Rabbit 2Y4824 anti-LPHN2 was diluted 1:300 for immunofluorescence analysis. 2Y4824 rabbit anti-pre-immune serum (PIS; Eurogentec) was used for control purposes. Mouse monoclonal (mAb) anti-HA tag (clone F-7) and goat polyclonal VE-Cadherin (clone C19) antibodies, used 1:200 for immunofluorescence analysis, and mAb anti-Shank2 (clone A11) used in immunoprecipitation, were from Santa-Cruz Biotechnology. Mouse mAbs anti-Vinculin (hVIN-1) and anti-α Tubulin (clone B-5-1-2) and rabbit polyclonal anti-Paxillin (HPA051309) were from Sigma-Aldrich and were diluted 1:400, 1:8000 and 1:200 respectively, for immunofluorescence and Western blot analyses. Rabbit polyclonal anti-ZO-1 (clone D6L1E) used in immunofluorescence 1:100 on cultured ECs was from Cell Signaling Technology. Mouse mAb anti-ZO-1 (clone ZO1-1A12) used in immunofluorescence 1:300 on zebrafish embryos was from Invitrogen. Rat mAb anti-HA High Affinity (clone 3F10), was from Roche and it was diluted 1:1000 for Western blot analysis. Mouse mAb anti-6xHis tag (clone HIS.H8) was from OriGene Technologies. Goat pAb anti-FLRT2 (AF2877), from R&D Systems, was diluted 1:100 and 1:1000 for immunofluorescence and Western blot analysis, respectively.

Horseradish peroxidase (HRP) conjugated donkey anti-goat (sc-2056), goat anti-rabbit (sc-2054) and goat anti-rat secondary Abs were from Santa Cruz Biotechnology, while HRP-goat anti-mouse (115-035-003) was from Jackson ImmunoResearch Laboratories. Alexa Fluor 555 donkey anti-mouse (A31570), Alexa Fluor 555 donkey anti-rabbit (A31572), Alexa Fluor 555 Phalloidin (A34055), Alexa Fluor 488 donkey anti-rabbit (A21206) and donkey anti-goat (A11055), Alexa Fluor 647 donkey anti-mouse (A31571), and DAPI (D3571) were from Thermo Fisher Scientific. Alkaline phosphatase (AP)-labeled goat anti-mouse IgG (H+L) cross-adsorbed secondary Ab (G-21060) was from Thermo Fisher Scientific.

### Recombinant proteins

Human fibronectin was from R&D Systems. Coll I from calf skin was produced by Sigma-Aldrich. Recombinant human carrier-free FLRT2 protein (2877-FL) was from R&D Systems. Streptavidin-conjugated HRP was from GE Healthcare.

### Gene silencing in cultured ECs

The day before oligofection, ECs were seeded in six-well plates at a concentration of 12 X 10^4^ cells per well. Oligofection of siRNA duplexes was performed according to the manufacturer’s protocols. Briefly, human ECs were transfected twice (at 0 and 24 h) with 200 pmol of siGENOME Non-Targeting siRNA #1 (as control) or siGENOME SMART pools siRNA oligonucleotides (GE Healthcare Dharmacon), using Oligofectamine Transfection Reagent (Thermo Fisher Scientific). 24 h after the second oligofection, ECs were lysed or tested in functional assays. Knockdown of human LPHN2 was achieved through the siGENOME SMART pool siLPHN2 M-005651-02 (siLPHN2 #1, 5’-GGAAUGGGCUUGCAAAGUU-3’; siLPHN2 #2, 5’-GGCAAGAACUCAAUUGAUU-3’; siLPHN2 #3, 5’-GAAAGAACGAGGAAUAUUG-3’; siLPHN2 #4, 5’-CGACAAACGUGCCGCAUCA-3’), while, in the case of human FLRT2, was the siGENOME SMART pool siFLRT2 M-009104-00 (siFLRT2 #1, 5’-GCAACCAACUGGACGAAUU-3’; si FLRT2 #2, 5’-CAGAUCGUCUCCUUAAAUA-3’; siFLRT2 #3, 5’-GGAACUUUGUCUACUGUAA-3’; siFLRT2 #4, 5’-UCAAAUAUAUCCCUUCAUC-3’). For silencing and rescue experiments, endogenous LPHN2 was silenced by transfecting the human-specific siGENOME SMART pool siLPHN2 M-005651-02 in ECs previously transduced with pCCL lentiviral vectors carrying or not (CTL) silencing resistant (see **Fig. S1E**) mouse WT or ΔOLF Lphn2 constructs.

### Western Blot analysis

Cells were lysed in buffer containing 25 mM Tris–HCl pH 7.4, 100 mM NaCl, 5% glycerol, 0.5 mM EGTA, 2 mM MgCl_2_, 1 mM PMSF, 1 mM Na_3_VO_4_, protease inhibitor cocktail (Sigma Aldrich) and detergent 1% NP40. Cellular lysates were incubated for 20 min on ice, and then centrifuged at 15,000 *g*, 20 min, at 4 °C. The total protein amount was determined using the bicinchoninic acid (BCA) protein assay reagent (Thermo Fisher Scientific). Specific amounts of protein were separated by SDS–PAGE with precast Bolt 4-12% Bis-Tris gel (Thermo Fisher Scientific) or Mini PROTEAN TGX precast 7,5% gel (Bio-Rad). Proteins were then transferred to a nitrocellulose membrane (Bio-Rad), probed with Abs of interest, and detected by enhanced chemiluminescence technique (PerkinElmer).

### Immunoprecipitation

To immunoprecipitate and analyze by Western blot endogenous LPHN2, 5 x 10^6^ HUVEC were transferred to ice, washed 3 times in cold PBS 1X, and surface-labeled at 4 °C with 0,2 mg/ml sulfo-NHS-SS-biotin (Thermo Scientific) in PBS1X for 30 min. Cells were then washed 3 times and lysed with a buffer containing 25 mM Tris–HCl pH 7.4, 100 mM NaCl, 1% Triton X-100, 5% glycerol, 0.5 mM EGTA, 2 mM MgCl_2_, 1 mM PMSF, 1 mM Na_3_VO_4_, and protease inhibitor cocktail (Sigma Aldrich). Cellular lysates were incubated for 20 min on ice and then centrifuged at 15,000 *g*, 20 min, at 4 °C. The total protein amount was determined using the bicinchoninic acid (BCA) protein assay reagent (Thermo Fisher Scientific). Equivalent amounts (1 mg) of protein were precipitated for 1 h at 4 °C with protein A-Sepharose beads and then centrifuge for 1 min 15,000 *g* at 4 °C. Protein A-Sepharose beads were collected to prepare the pre-clearing samples and the lysates were then immunoprecipitated for 1 h at 4 °C with the rabbit pAb anti-LPHN2 or pre-immune serum. Immunoprecipitates were washed four times with lysis buffer and then separated by SDS–PAGE. Proteins were then transferred to a nitrocellulose membrane (Bio-Rad), probed with Abs of interest, and detected by enhanced chemiluminescence technique (PerkinElmer).

### Rap1 and Rho-GTP pull-down assay

Rap1-GTP and Rho-GTP were respectively analyzed by means of the active Rap1 (Cat. #16120, Thermo Fisher Scientific) and Rho (Cat. #16116, Thermo Fisher Scientific) pull-down and detection kits. The assays were performed according to manufacturer’s protocols. Total Rap1 or Rho proteins, detected in the input fractions, were used to calculate the normalized amount of active Rap1-GTP or active Rho-GTP. Bands were quantified with ImageJ software and normalized optical density (N.O.D.) was calculated relative to control.

### Confocal microscopy on cultured ECs

Cells were plated on 0.17 mm glass coverslips (no. 1.5) pre-coated with FN [5 μg/ml] and allowed to adhere for 3 h. Cells were washed in phosphate-buffered saline (PBS), fixed in 4% or 2% paraformaldehyde (PFA), permeabilized in 0.1% Triton or 0,01% Saponin PBS 1X for 2 min or 5 min on ice respectively, incubated with different primary Abs and revealed by appropriate Alexa-Fluor-tagged secondary Abs (Thermo Fisher Scientific). Slides were mounted in Fluoromount-G (Southernbiotech, Cat. # 0100-01). Cells were analyzed at room temperature by using a Leica TCS SP8 AOBS confocal microscope equipped with two hybrid detectors (HyD) that, by combining classical photomultipliers with highly sensitive avalanche photodiodes, provides higher signal-to-noise ratio, image contrast and sensitivity. PL APO 63 x/1.4 NA immersion objective was employed. 1024×1024 pixel images were acquired, and a z-stack was acquired. For nucleus staining in immunofluorescence images taken with the Leica TCS SP8, TO-PRO-3 Fluorescent Nuclear Stain (ThermoFisher) was used. The acquisition was performed by adopting a laser power, gain, and offset settings that allowed maintaining pixel intensities (gray scale) within the 0–255 range and hence avoid saturation. Analysis of confocal images was performed using the ImageJ software package. FA and stress fiber number, FA size and stress fiber intensity were measured applying for each cell a default threshold. FA and stress fibers number was corrected on cell area. FA size was expressed as maximum Feret diameter. The fluorescence intensity of F-actin has been calculated adding the mean gray value of all the measured stress fiber within a cell.

To acquire super-resolved images, a Leica TCS SP8 gated-stimulated emission depletion (g-STED) 3X laser-scanning microscope equipped with a HC PL APO 100X/1.40 objective was employed (Leica Microsystems). Alexa Fluor 555 fluorochrome was excited at the optimal wavelength by means of 80 MHz pulsed white light laser (470–670 nm), allowing time gating of fluorescence lifetimes. For STED 660 nm depletion laser was used and emission was revealed by means of hybrid spectral detectors (HyD SP Leica Microsystems). Pixel size was maintained equal in all images. STED images were deconvolved to reduce noise using the mathematical algorithm Classic Maximum Likelihood Estimation (CMLE) included in Huygens Deconvolution Software. Stress fiber sagittal section and focal adhesion area was measured using ImageJ software.

Nuclear/cytoplasmic YAP and TAZ ratios were calculated by measuring the mean intensity fluorescence of each protein coming from the nucleus, divided by its cytoplasmic localization (cell area – nuclear signal).

ZO-1 localization to TJs was evaluated as percentage of VE-cadherin^+^ intercellular contact areas covered by ZO-1 staining.

### Binding assays

Empty vector, WT Lphn2 and ΔOLF Lphn2 expression constructs were transfected in COS-7cells to be used in the binding assay. *In situ* binding assays were performed as described previously (Tamagnone et al., 1999). Briefly, Lphn2-expressing COS-7 cells were seeded in wells of 48-well cluster dishes. They were then incubated for 1h at 37° with or without recombinant 6xHis-tagged human FLRT2, mouse mAb anti-6xHis, and AP-labeled goat anti-mouse. After five washes in DMEM diluted 1:2 in PBS, cells were fixed, heated for 10 min at 65°C to inactivate endogenous phosphatases, and incubated with NBT–BCIP (nitro blue tetrazolium–5-bromo-4-chloro-3-indolyl-phosphate) AP substrate (Promega, Catalog # S3771) for in situ cell staining. For a quantitative assessment of ligand binding, receptor-expressing cells were incubated with increasing concentrations of AP-conjugated ligands (with predetermined specific activity/µg); cell-bound AP activity was eventually revealed by incubation with the chromogenic soluble substrate p-nitrophenylphosphate (Sigma-Aldrich, Catalog # P7998) and measured by a multi-well spectrophotometer (absorbance at 405 nm).

### Endothelial cell migration assays

Real time directional EC migration was monitored with an xCELLigence RTCA DP instrument (ACEA Biosciences/Agilent Technologies) as previously described (Camillo et al., 2017; Gioelli et al., 2018). In details, the bottom side of the upper chamber (the side facing the lower chamber) of CIM-Plate 16 was coated with 30 μl of 1 μg/ml Coll I or 3 μg/ml FN for 30 min at room temperature. Each lower chamber well was first filled with 160 μl of M199 1% FBS (containing or not 800 ng/ml of rhFLRT2 protein) and then assembled to the upper chamber. Each upper chamber well was then filled with 30 μl of M199 1% FBS. The plate was put for 1h at 37°C. The experiment file was set up using the RTCA Software 1.2. ECs were detached and resuspended to a final concentration of 30,000 cells/100 μl. The BLANK step was started to measure the background impedance of cell culture medium, which was then used as reference impedance for calculating CI values. 100 μl of cell suspension (30,000 cells) were then added to each well of the upper chamber. The CIM-Plate 16 was placed in the RTCA DP Instrument equilibrated in a CO_2_ incubator. ECs migration was continuously monitored using the RTCA DP instrument. Average, standard deviation and p value were calculated on the CI data exported from RTCA instrument for the technical replicates of each experimental condition in the time. Migration data are represented as a percentage considering the control samples as 100%.

### cAMP assay

The cellular amounts of cAMP were quantified by using the non-acetylated cyclic AMP competitive ELISA Kit (Thermo Scientific, Cat. #EMSCAMPL) following to the manufacturer’s instructions. The concentration of cAMP in samples was expressed as pmol of cAMP per mg of total proteins.

### mRNA isolation from cultured ECs and real-time RT-PCR analysis

Cells were washed three times with PBS and frozen at −80°C. RNA isolation and reverse transcription: cells were thawed on ice, total RNA was extracted following the manufacturer’s recommended protocol (ReliaPrep RNA Miniprep Systems, Promega. Cat. # Z6011). The quality and integrity of the total RNA were quantified by the NanoDrop 1000 spectrophotometer (Thermo Scientific). cDNAs were generated from 1 µg of total RNA using the High Capacity cDNA Reverse Transcription Kit (Applied Biosystems). TaqMan real-time RT-PCR assay: mRNA expression of *LPHN2*, *FLRT1-3*, and endogenous housekeeping control genes, *i.e.* glyceraldehyde 3-phosphate dehydrogenase (*GAPDH*), and TATA binding protein (*TBP*), was measured by real-time RT-PCR using TaqMan Gene Expression Assays (Applied Biosystems) run on a C1000 Touch thermal cycler (Bio-Rad). The following assays were used: Hs00202347_m1 (*LPHN2*), Hs00534771_s1 (*FLRT1*), Hs00544171_s1 (*FLRT2*), Hs01922255_s1 (*FLRT3*), Hs00170014_m1 (*CTGF*), Hs00155479_m1 (*CYR61*), Hs99999905_m1 (*GAPDH*), and Hs00427620_m1 (*TBP*). For each sample, three technical replicates of each gene were run in a 96-well plate (cDNA concentration 50 ng/well) according the manufacturer’s protocol. Between the two measured housekeeping genes, we chose a normalization factor calculation based on the geometric mean of *GAPDH* and *TBP* gene transcript for most of real-time RT-PCR experiments, while *GAPDH* only was employed in **Fig. S2A** (Vandesompele et al., 2002). The experimental threshold (Ct) was calculated using the algorithm provided by the Bio-Rad CFX Manager 3.1 software (Bio-Rad). Ct values were converted into relative quantities using the method described by Pfaffl (Pfaffl, 2001) and amplification efficiency of each gene was calculated using a dilution curve and the slope calculation method (Pfaffl, 2001).

### Zebrafish embryo lines and handling

Zebrafish were handled according to established protocols and maintained under standard laboratory conditions. Experimental procedures related to fish manipulation followed previously reported recommendations (Workman et al., 2010) and conformed to the Italian regulations for protecting animals used in research, including DL 116/92. The Ethics committee of the University of Torino approved this study. Larvae were anesthetized and then sacrificed by ice chilling. Following zebrafish lines were used for these studies: wild-type AB, *Tg(kdrl:EGFP)^s843^*, carrying the endothelial-specific expression of the EGFP, and *Tg(Hsa*.*CTGF:nlsmCherry)^ia49^*/*Tg(kdrl:EGFP)^s843^*.

### Generation of *lphn2a* null zebrafish embryos

*Lphn2a^-/-^* zebrafish mutants were generated by CRISPR/Cas9-mediated genome editing at ZeClinics (Barcelona, Spain). A single guide RNA (sgRNA) was designed using the online tool http://crispor.tefor.net/, based on exon site, high efficacy, and not off-target published algorithms to specifically target an optimal CRISPR sequence on exon 2 of *lphn2a* gene (ENSDARG00000069356). The *lphn2a*-targeting sgRNA, with the specific targeting sequence CAACCGTCAAGACGAATACA-AGG, was injected in one-cell stage embryos in a solution containing Nls-CAS9 protein (PNA BIO). The mutagenesis efficacy was evaluated on pools of 30 injected embryos, whose genomic DNA was PCR amplified and analyzed via T7 endonuclease system. F0 injected embryos were raised to adulthood and screened, by genotyping the F1, for germline transmission of the mutation. Heterozygous mutants harboring the mutation were then incrossed to obtain homozygous mutants (F5 generation). The genomic region surrounding the CRISPR target site was PCR-amplified using the following primers: *lphn2a*-Fw (5’-TCTCAGAGTGACTTCCCCGGATC-3’), *lphn2a*-Rv (5’-GCAGCCATTATTTATCCCAGCTACC-3’). *Lphn2a^-/-^* heterozygous and homozygous mutants were identified by analyzing in agarose gel the PCR product digested with TasI restriction enzyme (TSP509I, Thermo Fisher Scientific) and by sequencing.

### Transmission electron microscopy (TEM)

Zebrafish embryos were fixed with 2.5% glutaraldehyde and 2% paraformaldehyde in 0.1M sodium cacodylate buffer pH 7.4 overnight at 4°C, following a standard TEM sample preparation protocol (Santoro et al., 2009). Briefly, the samples were postfixed with 1% osmium tetroxide in 0.1 M sodium cacodylate buffer for 2 hour at 4° C. After three water washes, samples were dehydrated in a graded ethanol series and embedded in an epoxy resin (Sigma-Aldrich). Ultrathin sections (60-70 nm) were obtained with an Ultrotome V (LKB) ultramicrotome, counterstained with uranyl acetate and lead citrate and viewed with a Tecnai G^2^ (FEI) transmission electron microscope operating at 100 kV. Images were captured with a Veleta (Olympus Soft Imaging System) digital camera.

### Larvae dissociation and fluorescence activated cell sorting (FACS)

WT and *Tg(kdrl:EGFP)^s843^* larvae at 2 days post-fertilization (dpf) were dissociated as previously described (Zancan et al., 2015) using 1x PBS, 0.25% trypsin phenol red free, 1 mM EDTA pH 8.0, 2.2 mg/ml Collagenase P (Sigma). Digestion was stopped by adding CaCl_2_ to a final concentration of 1 mM and fetal calf serum to 10%. Dissociated cells were rinsed once in PBS and resuspended in Opti-MEM (Gibco), 1% fetal calf serum and 1X Penicillin-Streptomycin solution (Sigma). Cells were filtered through a 40 μm nylon membrane. For sorting, we used FACS Aria IIIu sorter (BD Biosciences, San José, USA) with the following settings for EGFP: argon-ion Innova Laser (Coherent, USA) (488 nm, 100 mW); 100 μM nozzle; sorting speed 500 events/s in 0-32-0 sort precision mode. We performed data acquisition and analysis with the BD FACSDiva software (BD Biosciences, San José, CA, USA). GFP^+^ and GFP^-^ cells were separately collected in resuspension medium, and RNA was extracted using the RNA isolation kit Nucleospin_®_ RNA XS (Macherey-Nagel). cDNA was made with RT High Capacity kit (Applied Biosystems) according to the manufacturer’s protocol. Quantitative real-time RT-PCR on ECs from zebrafish was performed with CFX384 Touch Real-time PCR Detection System (Biorad) using 5x HOT FIREPol^®^EvaGreen^®^ qPCR Mix Plus (Solis BioDyne), the following primer sequences were used *lphn2*-F: 5’-AGTATCCCTCATCTGCCTGG-3’ *lphn2*-R: 5’-AGCTGAACTCCTTCCAGACA-3’; *flrt2*-F: CATTGCATGGCTCAGGTCTC, *flrt2*-R: ATGAGTTGGCCAGGGATGAA, *cyr61*-F: 5’-GCGGAGACTCGGAGAAAGAAC-3’, *cyr61*-R 5’-CGATGCACTTCTCCATCTGATG3’; *ctgfa*-F: 5’-CTCCCCAAGTAACCGTCGTA-3’, *ctgfa*-R: 5’-CTACAGCACCGTCCAGACAC-3; *ctgfb*-F: 5’-CCCACAAGAAGACACCTTCC-3’, *ctgfb*-R: 5’-ATTCGCTCCATTCAGTGGTC-3’. Results are expressed as relative mRNA abundance and normalized to actin beta1 (*actb1*) or eukaryotic translation elongation factor 1 alpha 1, like 1 (*eef1a1l1*) as endogenous reference genes, which were amplified by employing the following primer sequences *actb1*-F: 5’ GTATCCACGAGACCACCTTC-3’, *actb1*-R: 5’-GAGGAGGGCAAAGTGGTAAAC-3’; *eef1a1l1*-F 5’-GACAAGAGAACCATCGAG-3’, *eef1a1l1*-R 5’-CCTCAAACTCACCGACAC-3’.

### Assessment of vascular permeability in zebrafish embryos

*Lphn2a*^+/-^ fish were incrossed and the progeny were incubated in 0.003% 1-phenyl-2-thiourea (PTU) to inhibit pigment formation. The vascular permeability experiment was performed as previously described (Hoeppner et al., 2012). Microangiography was performed on anesthetized 3 days post fertilization (dpf) embryos, by injecting in the duct of Cuvier a solution containing FITC-dextran (70 kDa) (Life Technologies, Inc.) at 1 mg/ml concentration. Vascular Endothelial Growth Factor human (VEGF Sigma, Cat. # V7259) was injected in the duct of Cuvier. The visualization and real-time imaging were performed after 3 hours on a Leica SP8 confocal microscope. The average of the dextran fluorescence in the intervascular areas was normalized to the average dextran fluorescence inside the vessels and was quantified after the embryos genotyping.

### Intravascular injection of cancer cells in zebrafish embryos

At 48 hpf, embryos were dechorionated, anesthetized with tricaine 0.16 mg/ml and placed along plastic lanes immersed in 2% methylcellulose/PBS. Sub-confluent cells (B16F10 or SK-MEL-28) in 10 cm culture dishes were rinsed twice with warm serum free medium and then incubated for 30 minutes at 37°C 200nM with MemBright-560 (Lipilight-IDYLLE). To eliminate all possible traces of unbound MemBright-560, cells were rinsed three times with serum free medium, and then cells were harvested using trypsin-EDTA solution. Stained cells (at 100 · 10^6^ cells per ml) were loaded in a glass capillary needle and microinjected into the duct of Cuvier of the embryos under the stereomicroscope using a WPI PicoPump apparatus. Xenotransplanted embryos were grown at 32° C, monitored daily, and analyzed starting from one day post injection (dpi) up to 3 days.

### Image acquisition and analysis on zebrafish embryos

Double transgenic fluorescence of *Tg(kdrl:EGFP)^s843^* and Tg *(Hsa.CTGF:nlsmCherry)* in *lphn2a*^-/-^ mutant background was visualized under a AZ100 stereomicroscope equipped with AxioCam (Zeiss) dissecting microscope and then with a Leica SP8 confocal microscope at room temperature. HC PL APO 20x/0.75 IMM CORR CS2 objective was employed. Larvae were anaesthetized and mounted in 1% low-melting-point agarose 1,5% gel. Endothelial cells targeted EGFP and nuclear mCherry fluorescence (Yap/Taz) was visualized by using 488 nm and 561 nm lasers. All images were analyzed with the *3D ImageJ Suite* of ImageJ/Fiji (Ollion et al., 2013). The 3D analysis of Figs. 4C and 4E was performed using the Gaussian Blur 3D filters followed by the 3D simple segmentation using methods of the 3D ImageJ Suite. Yap/Taz signal were automatically segmented on the mCherry fluorescence image stacks, inside a EGFP fluorescent mask identifying endothelial cells, and total signal intensity was calculated, according to previous work (Facchinello et al., 2016). After confocal acquisition, heterozygous and homozygous siblings were genotyped by PCR on DNA previously extracted from a single larva, as described by (Gagnon et al., 2014).

### Statistical analysis

For statistical evaluation of *in vitro* experiments, data distribution was assumed to be normal, but this was not formally tested. Parametric two-tailed heteroscedastic Student’s t-test was used to assess the statistical significance when two groups of unpaired normally distributed values were compared; when more than two groups were compared, parametric two-tailed analysis of variance (ANOVA) with Bonferroni’s post hoc analysis was applied. For all quantifications, Standard Deviation (SD) is shown. All data were analyzed with Prism software (GraphPad Software, San Diego, CA).

For statistical evaluation of *in vivo* experiments, neither randomization nor blinding was applied for samples or zebrafish embryo analyses. No statistical method or criteria was used to predetermine sample size or to include/exclude samples or animals. The Shapiro-Wilk normality test was used to confirm the normality of the data. The statistical difference of Gaussian data sets was analyzed using the one-way ANOVA with Tukey’s multiple comparison test, in case of unequal variances. For data not following a Gaussian distribution, the Mann-Whitney test was used. Illustrations of statistical analyses of *in vivo* experiments are presented as the means ± SD.

For both *in vitro* and *in vivo* analysis, statistical differences were considered not significant (NS) = p-value > 0,05; significant * = p-value ≤ 0,05; ** = p-value ≤ 0,01; *** = p-value ≤ 0,001).

Supplementary Figs. 1, 2, and 3 are available online. Raw data of all graphs and uncropped scans of Western blots are publicly available on Figshare public repository (https://doi.org/10.6084/m9.figshare.15164292).

## ACKNOWLEDGEMENTS

We are grateful to Prof. Alberto Puliafito (University of Torino) for help with data analysis and to Prof. Georg Halder (University of Leuven) for fruitful discussion on YAP/TAZ and cell adhesion. We thank the zebrafish facility at UNIPD. The research leading to these results has received funding from: Fondazione AIRC IG grants #13016, #16702, and #21315 (to G.S.), #19923 (to L.T.), #20366 (to D.V.), and #20119 (to M.M.S.); Fondazione AIRC under 5 per Mille 2018 - ID. 21052 program – P.I. Comoglio Paolo, G.L. Tamagnone Luca, G.L. Serini Guido; FPRC-ONLUS Grant “MIUR 2010 Vaschetto - 5 per mille 2010 MIUR” (to G.S.); Telethon Italy (GGP09175) (to G.S.); Università di Torino, Bando Ricerca Locale 2019 (CUP D84I19002940005) (to G.S.); Associazione ‘Augusto per la Vita’ (to G.S.); ERC-CoN project 647057 - rEnDOx (to M.M.S.). The authors declare no further competing financial interests.

## AUTHOR CONTRIBUTIONS

G Serini conceived the project; G Serini, C Camillo, G Villari, G Mana, D Valdembri, MM Santoro, and N Facchinello designed the experiments; G Serini, G Villari, C Sandri, D Valdembri, and MM Santoro supervised the research; L Tamagnone and M Arese provided key reagents, methods, and technologies; C Camillo, N Facchinello, G Villari, G Mana, C Sandri, RE Oberkersch, D Tortarolo, F Clapero, D Gays, and N Gioelli performed the experiments; C Camillo, N Facchinello, G Villari, G Mana, C Sandri, RE Oberkersch, D Tortarolo, M Arese, F Clapero, D Gays, and N Gioelli, D Valdembri, M Arese, L Tamagnone, MM Santoro, and G Serini analyzed the data; C Camillo, N Facchinello, G Villari, G Mana, C Sandri, RE Oberkersch, D Tortarolo, M Arese, F Clapero, D Gays, and N Gioelli, D Valdembri, M Arese, L Tamagnone, MM Santoro, and G Serini interpreted the results; C Camillo, N Facchinello, G Villari, D Valdembri, MM Santoro, and G Serini wrote the paper; all authors read and approved the manuscript.

---

**Figure S1.**
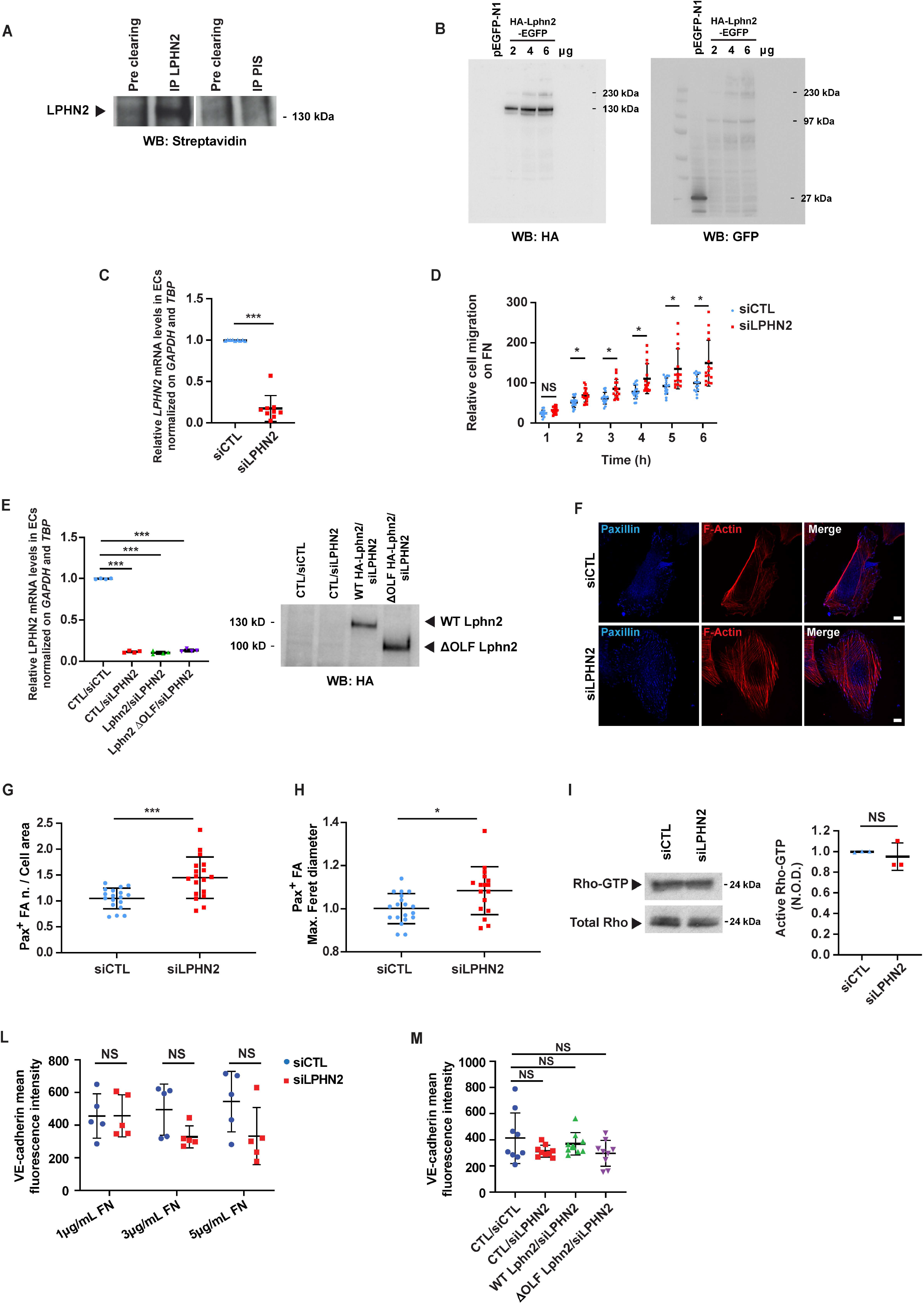
(**A**) Upon surface biotinylation LPHN2 was immunoprecipitated with a rabbit anti-LPHN2 pAb from EC lysates and revealed in Western blot by means of HRP streptavidin. Pre-immune serum (PIS) was employed for control purposes. The cleaved extracellular portion of LPHN2 appears as a ∼130 kDa protein band. (**B**) HEK 293T cells were transfected with an empty vector or increasing amounts (2, 4 and 6 μg) of HA-Lphn2-EGFP construct. Cells lysates were then analyzed by Western blot with either anti-HA (left panel) or anti-GFP (right panel) antibody. The 230 kDa, 130 kDa, and 97 kDa protein bands correspond to uncleaved full-length, cleaved N-terminal extracellular-only portion, and cleaved C-terminal portion of the transfected HA-Lphn2-EGFP protein, respectively. In the anti-GFP Western blot, the 27 kDa band corresponds to the GFP whose cDNA was present in the control empty vector. (**C**) Real time quantitative PCR analysis of *LPHN2* mRNA in siCTL or siLPHN2 human ECs relative to the house keeping genes *GAPDH* and *TBP* and normalized on siCTL levels. Results are the average ± SD of nine independent assays. Statistical analysis: two-tailed heteroscedastic Student’s t-test; p≤ 0,001 ***. (**D**) Real time analysis of control (siCTL) or LPHN2 (siLPHN2) silenced EC migration toward FN was assessed with an xCELLigence RTCA DP system. Results are the average ± SD of four independent assays. Statistical analysis: two-way ANOVA and Bonferroni’s post hoc analysis; p> 0,05 NS; p≤ 0,05 *. (**E**) Real time quantitative PCR analysis of *LPHN2* mRNA in siCTL or siLPHN2 human ECs rescued or not (CTL) with mouse WT or ΔOLF Lphn2 relative to the house keeping genes *GAPDH* and *TBP* and normalized on siCTL levels. Results are the average ± SD of four independent assays. Statistical analysis: one-way ANOVA and Bonferroni’s post hoc analysis; p ≤ 0,001 ***. Right panel, transduced and silenced ECs were also lysed and WT HA-Lphn2-pCCL and ΔOLF HA-Lphn2-pCCL protein expression levels analyzed by Western blot with anti-HA antibody. (**F-H**) Confocal microscopy analysis (**F**) of endogenous paxillin (*blue*), and phalloidin-labeled F-actin (*red*) reveals how, compared to siCTL ECs, LPHN2 silencing increases the number, normalized on cell area (**G**) and size (expressed by maximum Feret diameter **H**) of paxillin-containing FAs. Scale bars: 10 µm. Results are the average ± SD of two independent experiments for a total of 18 (siCTL) and 18 (siLPHN2) ECs. Statistical analysis: two-tailed heteroscedastic Student’s t-test; p≤ 0,05 *; p <0,001 ***. (**I**) LPHN2 silencing in human ECs does not affect basal GTP loading of RhoA small GTPase. Total Rho was used to calculate the normalized optical density (N.O.D.) levels of active Rho-GTP. Results are the average ± SD of three independent assays. Statistical analysis: two-tailed heteroscedastic Student’s t-test; p> 0,05 NS. (**L**) Mean fluorescence intensity of VE-cad^+^ intercellular staining (in *green* in Fig. 5A) in siCTL ECs seeded on 10 kPa substrates coated with increasing amounts of FN (1, 3, and 5 µg/ml). The VE-cadherin intercellular recruitment was not affected by FN density. Results are the average ± SD of two independent experiments. Statistical analysis: two-way ANOVA with Bonferroni’s post hoc analysis; p> 0,05 NS. (**M**). Mean fluorescence intensity of VE-cad^+^ intercellular staining (in *green* in Fig. 5B) in lentivirally-delivered WT or ΔOLF Lphn2 in ECs seeded on 10 kPa substrates coated FN (5 µg/ml)-coated coverslips. The VE-cadherin intercellular recruitment was not affected by LPHN2 silencing. Results are the average ± SD of two independent experiments. Statistical analysis: two-way ANOVA with Bonferroni’s post hoc analysis; p> 0,05 NS.

**Figure S2.**
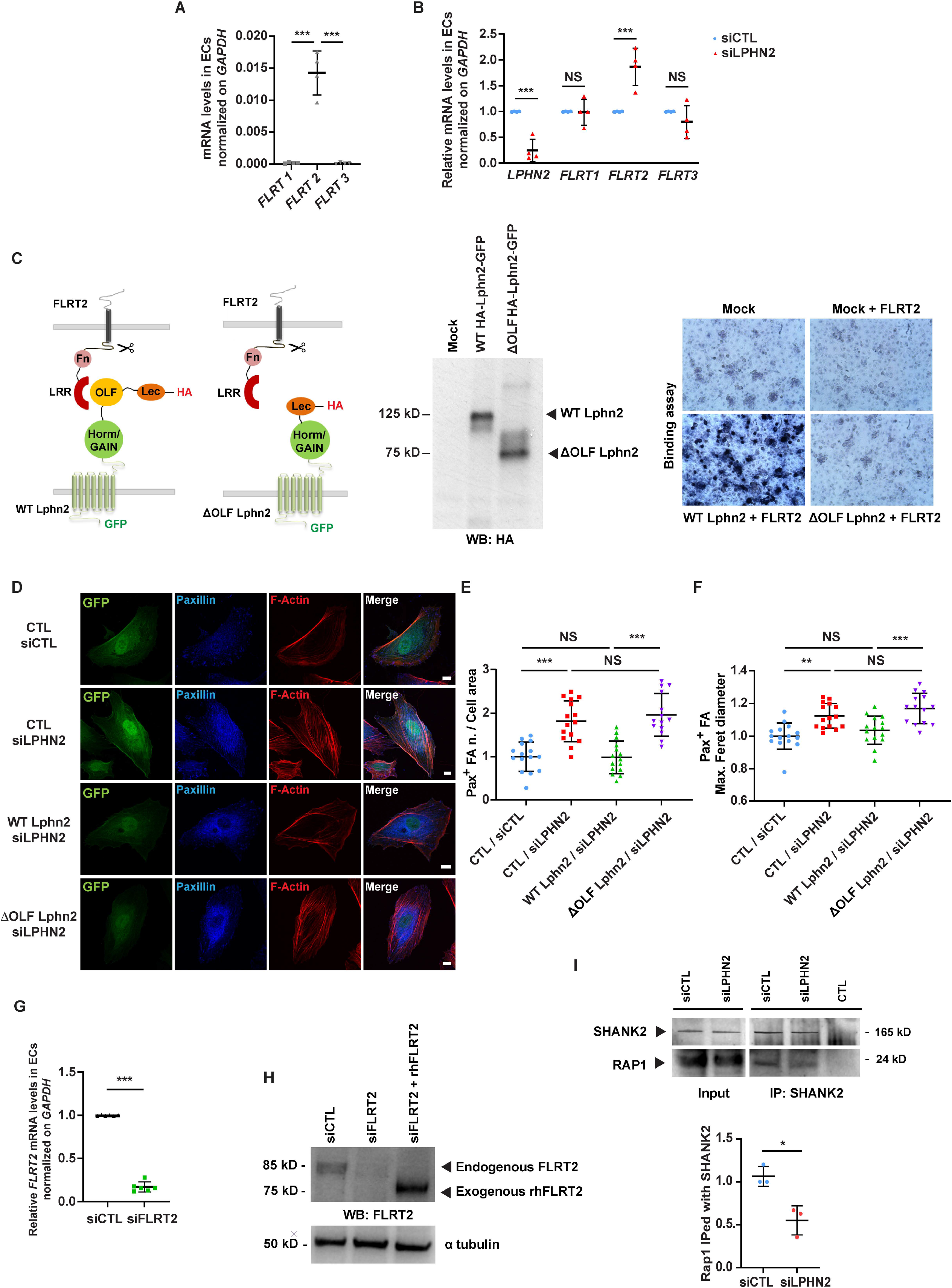
(**A**) Real time quantitative PCR analysis of *FLRT1-3* mRNAs in human ECs relative to the house keeping gene *GAPDH*. Results are the average ± SD of four independent experiments. Statistical analysis: one-way ANOVA and Bonferroni’s post hoc analysis; p ≤ 0,001 ***. (**B**) Real time quantitative PCR analysis of *LPHN2* and *FLRT1-3* mRNAs in siCTL and siLPHN2 human ECs. Results are normalized on siCTL values and are the average ± SD of four independent experiments. Statistical analysis: one-way ANOVA and Bonferroni’s post hoc analysis; p> 0,05 NS; p≤ 0,001 ***. (**C**) COS-7 cells transfected with empty vector or WT HA-Lphn2-EGFP or ΔOLF HA-Lphn2-EGFP constructs and their expression verified by Western blot analysis employing a rat mAb anti-HA (middle). Next, COS-7 cells transfected with empty vector or WT HA-Lphn2-EGFP or ΔOLF HA-Lphn2-EGFP constructs were incubated or not with 6xHis-tagged rhFLRT2. The binding between the Lphn2 constructs and the rhFLRT2 ligand was revealed through sequential incubation with a mouse mAb anti-6xHis, an AP-conjugated goat anti-mouse pAb, and the AP substrate NBT–BCIP (right). Results show how WT, but not ΔOLF Lphn2 binds rhFLRT2. (**D-F**) Confocal microscopy analysis (**D**) of paxillin (*blue*), and phalloidin-labeled F-actin (*red*). Cells were first transduced with pCCL lentivirus (carrying GFP)-mediated overexpression (*green*) of silencing-resistant mouse WT or ΔOLF Lphn2 and then oligofected with either siCTL or siLPHN2 siRNAs. Scale bar: 10 µm. Confocal microscopy analysis reveals how, lentiviral delivery of WT Lphn2, but not ΔOLF Lphn2 mutant, restores the phenotype of paxillin-containing FAs both considering the number (**E**) and the size (expressed by maximum Feret diameter, **F**). Results are the average ± SD of 2 independent experiments for a total of 15 ECs for each condition. Statistical analysis: one-way ANOVA and Bonferroni’s post hoc analysis; p> 0,05 NS; p≤ 0,01 **; p≤ 0,001 ***. (**G**) Left panel, real time quantitative PCR analysis of *FLRT2* mRNA in siCTL or siFLRT2 human ECs relative to the housekeeping gene *GAPDH* and normalized on siCTL levels. Results are the average ± SD of six independent assays. Statistical analysis: two-tailed heteroscedastic Student’s t-test; p≤ 0.001 ***. Right panel, Western blot analysis with an anti-FLRT2 Ab of lysates of siCTL or siFLRT2 ECs. (**H**) Western blot analysis with an anti-FLRT2 Ab of lysates of siCTL or siFLRT2 ECs or siFLRT2 ECs treated with exogenous rhFLRT2 [800 ng/ml]. Endogenous FLRT2 appears as a ∼85 kDa protein, while the soluble extracellular portion of exogenous rhFLTR2 appears as a ∼75 kDa protein band. (**I**) LPHN2 silencing in human ECs decreased SHANK2 interaction with Rap1 small GTPase. Western Blot analysis of Rap1 co-immunoprecipitated with SHANK2 in cultured ECs. Results are the average ± SD of three independent experiments. Statistical analysis: two-tailed heteroscedastic Student’s t-test; p≤ 0,05 *.

**Figure S3.**
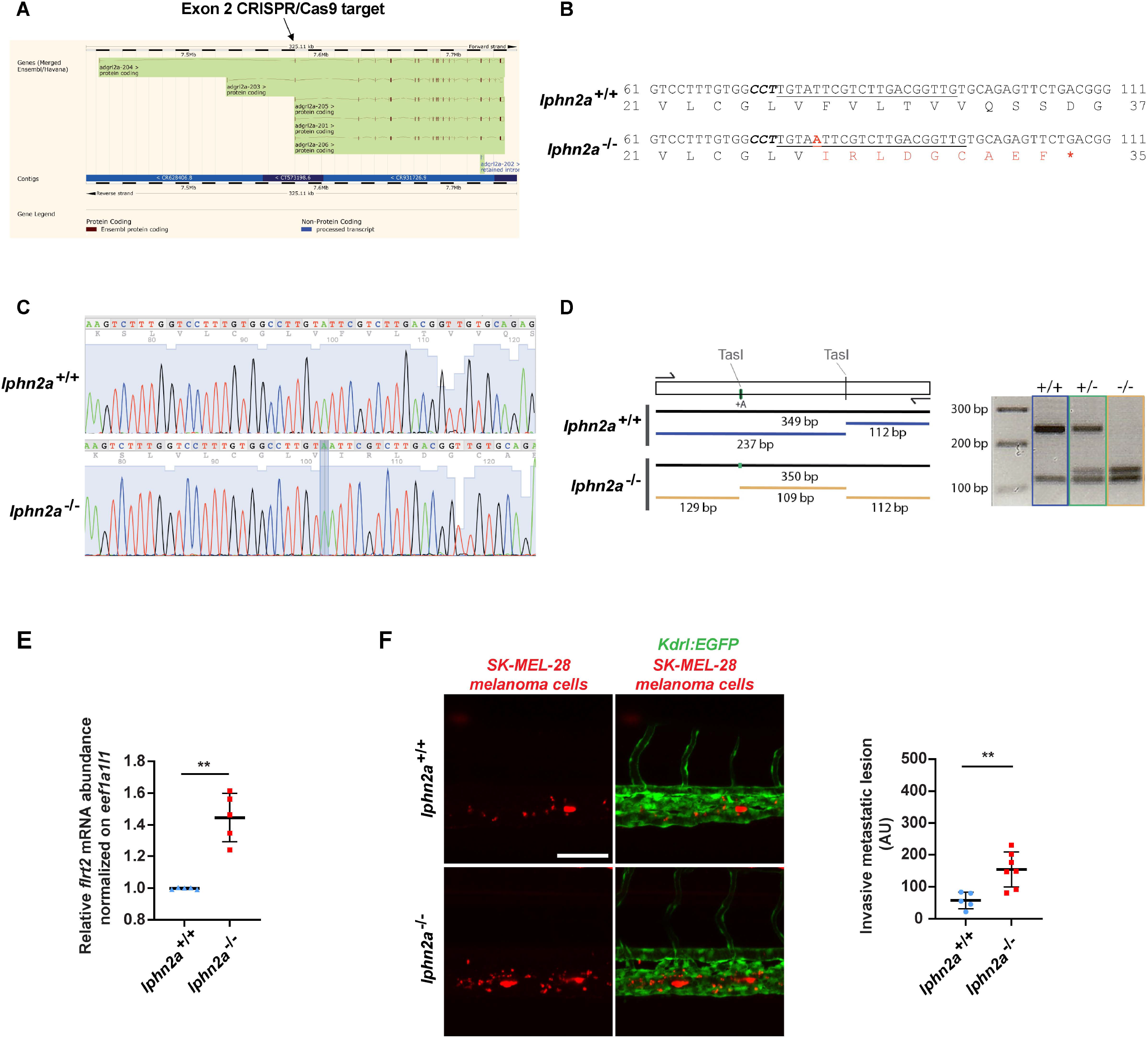
(**A**) *lphn2a* (aka *adgrl2a*) zebrafish mRNA splicing variants displaying all exons. The black arrow outlines the lphn2a gene targeted region located in the second exon, which is common to all splicing variants. (**B**) DNA sequence details of the exon 2 *lphn2a* gene targeted region in *lphn2a^+/+^* and *lphn2a^-/-^* zebrafish embryos. Locus specific sgRNA is underlined and the protospacer-adjacent motif (PAM) sequence is labelled in bold italic. A 1-nucleotide insertion (A, *red bold underscored*), revealed by sequencing, and the consequent nine new amino acids (*red*) and STOP codon (*red bold*) are shown. (**C**) Alignment of *lphn2a^+/+^* and *lphn2a^-/-^* sequence obtained by Sanger sequencing. (**D**) Schematic representation of genotyping. The one base insertion generates a second TasI restriction site in the diagnostic PCR, which has been used for genotyping. (**E**) Real time quantitative RT-PCR analysis of *flrt2* mRNA in *lphn2a^+/+^* or *lphn2a^-/-^* zebrafish embryos relative to the house keeping gene *eef1a1l1* and normalized on the mRNA levels measured in *lphn2a^+/+^* animals. Results are the average ± SD of 5 independent assays (n>80 embryos for condition). Statistical analysis: Mann-Whitney test; p≤ 0,01 **. (**F**) MemBright-560-labeled melanoma cells were microinjected into the duct of Cuvier of 48 hpf *lphn2a^+/+^* or *lphn2a^-/-^ Tg(Kdrl:EGFP)* zebrafish embryos. After 36 hours, extravasated metastatic melanoma cells were imaged by confocal analysis of the caudal plexus. Human SK-MEL-28 melanoma cell extravasation is enhanced in *lphn2a^-/-^* compared to *lphn2a^+/+^* zebrafish embryos. Results are the average ± SD of 2 independent assays, in which 12 SK-MEL-28 melanoma cell-injected animals were analyzed. Statistical analysis: Mann-Whitney test; p≤ 0,01 **.

